# Neural state space alignment for magnitude generalisation in humans and recurrent networks

**DOI:** 10.1101/2020.07.22.215541

**Authors:** Hannah Sheahan, Fabrice Luyckx, Stephanie Nelli, Clemens Teupe, Christopher Summerfield

**Affiliations:** Department of Experimental Psychology, University of Oxford, Oxford, UK

## Abstract

A prerequisite for intelligent behaviour is to understand how stimuli are related and to generalise this knowledge across contexts. Generalisation can be challenging when relational patterns are shared across contexts but exist on different physical scales. Here, we studied neural representations in humans and recurrent neural networks performing a magnitude comparison task, for which it was advantageous to generalise concepts of “more” or “less” between contexts. Using multivariate analysis of human brain signals and of neural network hidden unit activity, we observed that both systems developed parallel neural “number lines” for each context. In both model systems, these number state spaces were aligned in a way that explicitly facilitated generalisation of relational concepts (more and less). These findings suggest a previously overlooked role for neural normalisation in supporting transfer of a simple form of abstract relational knowledge (magnitude) in humans and machine learning systems.

## Introduction

Humans can think and reason in ways that abstract over the physical properties of the world (B. M. Lake et al., 2017; Tenenbaum et al., 2011). For example, we understand that cheetahs and space rockets can both move “fast” even though animals and vehicles belong in different semantic categories and do not look alike. Cognitive scientists have long built theories about how humans learn concepts and reason abstractly but much less is known about their neural representation (Gentner, 2010; Murphy, 2002). One view is that conceptual knowledge relies on neural ensembles that code for relations among stimuli but are invariant to their physical properties (Baram et al., 2019; Behrens et al., 2018; Bellmund et al., 2018; Collins & Frank, 2013; Doumas et al., 2008; Brenden M. Lake et al., 2015; Summerfield et al., 2019; Tervo et al., 2016). Recent evidence hints that when sets of stimuli share relational structure across contexts, they are embedded on parallel low-dimensional neural manifolds, so that a linear decoder learned in one context can be readily repurposed for another (Bernardi et al., 2020; Fitzgerald et al., 2013; Ganguli et al., 2008; Luyckx et al., 2019; Remington et al., 2018). By aligning neural state spaces between contexts in this way, one can generalise relational knowledge, for example applying a criterion that distinguishes fast and slow animals to discriminate fast and slow vehicles, such as space rockets and bicycles (**Fig. 1a**). The neural geometry implied by this coding scheme thus offers a theory for how humans engage in abstract forms of reasoning that involve the use of analogy and metaphor (Gentner, 2010).

**Fig 1.**
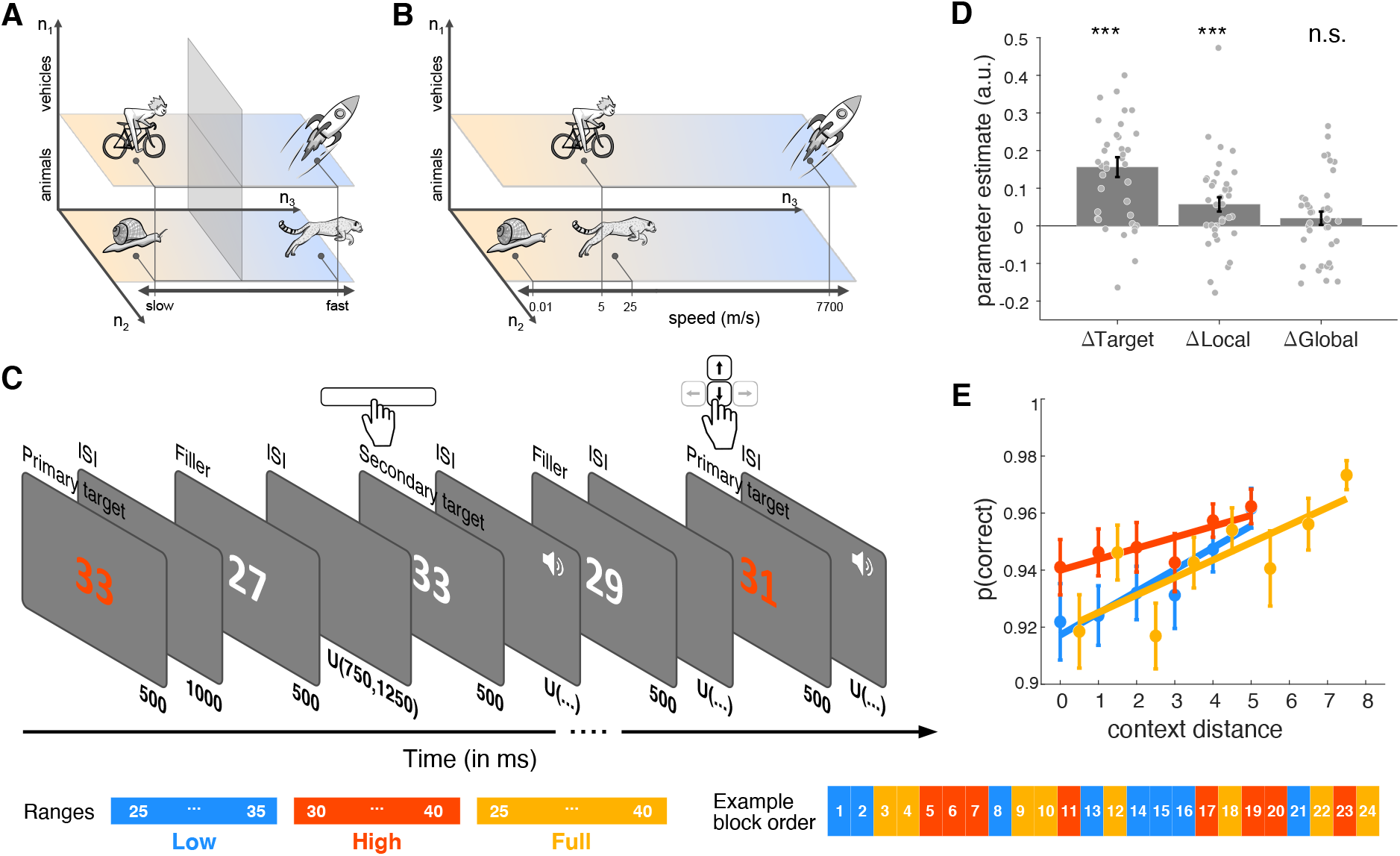
The mapping problem, the task and human behavioural results. (A) A sketch of aligned neural representations of stimuli in different contexts (e.g. animals, vehicles), in a 3-dimensional neural space spanned by the firing rates of 3 hypothetical neurons [n_1_, n_2_, n_3_]. One of these neurons encodes magnitude (n_3_) while another encodes context (n_1_). In this example, a decoder (grey vertical plane) trained to discriminate the relative speed of bicycles and space rockets could be repurposed to distinguish snails from cheetahs. Such decoder generalisation permits context-general inferences, such as saying that the space rocket and the cheetah are both “fast” even though space rockets and cheetahs belong in different categories and do not look alike. (B) In contrast, a coding scheme based on an absolute metric, such as numerical speed in m/s, positions stimuli that are in different contexts but with the same within-context relations, very differently. (C) Schematic of the task performed by humans. The primary task was to compare the relative magnitude of primary target numbers (coloured) and indicate a binary “more” or “less” response using the up/down keys. Numbers were drawn from three temporally blocked ranges (contexts): a low range (blue), a high range (red) and a full range (golden). Blocks of trials were presented eight times per context, and the order of blocks was pseudorandom. (D) Parameter estimates from logistic regression of human accuracy against the target distance (Δ Target), local context distance (Δ Local) and global context distance (Δ Global). Dots show individual subject fits. (E) Human accuracy increases as a function of context distance for each context. Error bars show s.e.m in panels D and E.

However, there is significant challenge for relational generalisation that we call the *mapping problem*. The mapping problem occurs when stimuli are analogously related across contexts, but in one context the structure is rotated, rescaled or otherwise misaligned with respect to the other. To illustrate, a spectator of Olympic events might consider both a record-breaking sprinter and a marathon champion to be “fast” runners, but in one case this might entail running one hundred metres in less than ten seconds, and in the other a marathon in under two hours (**Fig. 1b**). For generalisation to be effective, we need to form a concept of “fast” that is not tied to a specific physical value (e.g. a number denoting m/s), but that encodes relative speed in each respective context. Understanding how these abstract sorts of invariances are acquired in either biological or artificial neural networks, however, remains a challenge to researchers in neuroscience and machine learning alike.

Here, we use simulations with recurrent neural networks and electroencephalographic (EEG) recordings from humans to ask how stimuli with a common relational structure across contexts were represented in neural state spaces, with a focus on the mapping problem. We constructed a task which involved comparing the magnitude of arbitrary, physically dissimilar stimuli that were sampled from one of three overlapping ranges (contexts). For humans, we used symbolic numbers (Arabic digits) as stimuli because adults have already learned that they denote positions on a one-dimensional manifold (number line) with visual symbols that are physically dissimilar in arbitrary ways. For neural networks, we use arbitrary non-overlapping (one-hot) inputs whose assigned magnitude could be inferred via supervision signals during training. In the context of the Olympics example above, the number corresponds to a speed (e.g. in m/s) and the different contexts correspond to temporally segregated competitions (sprint vs. marathon), and our question is how the brain learns the concept of “fast” vs. “slow”. When we conducted multivariate analyses on neural signals or hidden unit activations observed during the task, using dimensionality reduction to visualise the structure of the neural state spaces, we observed neural state space alignment across contexts in both humans and neural networks. In both model systems, numbers are organised according to their magnitude onto three parallel, equidistant neural manifolds (number lines), one for each context. Moreover, in both systems these manifolds were compressed (divisively normalised) and centred (subtractively normalised) so that numbers that denoted “more” or “less” could be linearly discriminated along a single dimension irrespective of their context. In other words, without being explicitly regularised to do so, neural networks autonomously learned to align representations in a way that supports generalisation between contexts. In doing so, they learned to represent numbers with a neural geometry that matched that in the brains of human participants. This suggests a hitherto overlooked role for neural normalisation (Carandini & Heeger, 2012) in supporting learning and transfer of relational knowledge.

## Results

The task was a serial visual presentation paradigm which required agents (humans or neural networks) to classify the transitive ordering of two sequential target stimuli, indicating whether the current target was “higher” or “lower” than the previous target. For humans, stimuli were Arabic two-digit numbers and the correct transitive ordering was given by their numerical magnitude (so 21 was “lower” than 22). For neural networks, each stimulus corresponded to an arbitrary (or one-hot) input (equivalent to a number) provided to the network, and the transitive ordering was learned *tabula rasa* via supervision signals administered in a training phase (see below). So they network learned, after training, to respond “lower” when the current target inputs was that denoting 21 and the previous had been that denoting 22.

For both humans and neural networks, target magnitudes were drawn from different ranges in three temporally blocked contexts, which were repeated in pseudorandom order (24 blocks, 8 blocks in each context), and in each block participants viewed a sequence of 120 stimuli, of which 30 were primary targets. Target stimuli occurred in each of three contexts, which uniformly spanned digit ranges 25-35 (low), 30-40 (high) and 25-40 (full context) in distinct blocks. For humans, targets were signalled by a distinctive cue that was also indicative of the context (font colour of the number), and responses were made with two fingers of the right hand. Interspersed between successive targets were 2-4 “filler” numbers (white for humans) that were drawn from across the full range. A secondary task that did not require higher vs. lower magnitude comparison was performed on these stimuli by the humans, namely pressing the space bar (with the left hand) when a filler exactly matched the previous target (~12% of fillers). A visualisation of the task that humans performed is shown in **Fig. 1c**.

### Human behaviour

We first focus on the primary (magnitude comparison) task which healthy human adults (n = 38) performed with a mean accuracy of ~94±4% and response times that averaged 675ms. We begin by offering a normative account of this task. An observer with perfect memory can ignore the context provided by recent numbers and simply maintain the previous item for comparison with the current item (e.g. compare **31** with **33** in **Fig. 1c**). However, for an agent with imperfect memory, the context in which numbers occur becomes relevant. For example, a participant who forgets the **33** but responds “less” to **31** will most likely be correct in the high context (Holllingworth, 1910; Jazayeri & Shadlen, 2010). Critically however this latter strategy will be more effective when participants have some notion of whether each target is “more” or “less” relative to the local context within the current block, because a number may be low in one context but high in another (for example **31** is “more” than the local average in the low context, and so would prompt the incorrect answer). This effect is well known to lead to estimation biases that depend on the local history and have been associated with neural signals in the parietal cortex (Akrami et al., 2018).

Here, behavioural analyses revealed that participants used memory and contextual information when making decisions (**Fig. 1d**). Logistic regression indicated that accuracy was predicted by both the disparity between current and past target number (*target distance*; t_37_ = 5.9, p < 0.001) and the distance between the target number and the median number in the current block (*local context distance* t_37_ = 3.0, p < 0.004). After accounting for these sources of variance, however, distance to the overall median number (across all blocks; *global context distance*) had no impact on performance (p > 0.1). We plot accuracy as a function of local context distance for the low, high and full contexts in **Fig. 1e**. Under this normative account, context should have more influence when more time has gone by, under the assumption that target information becomes more fragile with longer delays. This follows naturally from the idea that when current information is weak or ambiguous, normative agents rely on the central tendency of experience (Akrami et al., 2018; Holllingworth, 1910; Jazayeri & Shadlen, 2010). We thus also compared the influence of context on accuracy and RT when there were either 2, 3 or 4 intervening filler by including an additional term in the previous regression to encode the interaction between number of fillers and the local context. Despite a tendency for this predictor to trend in the expected direction of context being more influential after delays that included more fillers, it was not significant for either accuracy (t_38_ = 1.16, p = 0.13, one-tailed) or RT (t_38_ = −1.56, p = 0.06, one-tailed).

### Neural network behaviour

For comparison, we trained recurrent neural network models (RNNs; n = 10) to perform an equivalent task on symbolic (one-hot) inputs (see **Fig. 2a**). As for human experiments, inputs were sampled from different ranges in blocks of 120 trials. Each network had a single recurrent layer and a single feedforward hidden layer and was trained to minimise errors on the task. Importantly, the RNN is not reset after every trial, but initialised at the start of the first block of training. The hidden state at the end of each block served as the initial hidden state for the subsequent block. We encouraged the network to find context relevant in the same way as humans using a virtual inactivation (VI) approach. This involved setting the input to zero for a fraction ε of targets, as if stimuli were missed or forgotten, as they might be by a human with imperfect memory. Applying VI during training obliged the network to learn to use the context (i.e. local average of numbers) as a cue for magnitude comparison, because when numbers were lost to memory, knowledge of the context improved accuracy. Similarly, many studies have shown that the influence of different sources of information to human decisions (such as the context, or the number) depends on the reliability of that source (Ernst & Banks, 2002; K. Kording, 2007; K. P. Kording & Wolpert, 2004). Applying VI during test allowed us to measure this impact of context on responding independent of the target distance. After ~10^6^ training steps, networks converged to near perfect accuracy on held out stimulus sequences irrespective of whether virtual inactivation was applied during training at ε_*train*_ = 0.1 (99.80 ± 0.05%) or not (99.96 ± 0.03%).

**Fig 2.**
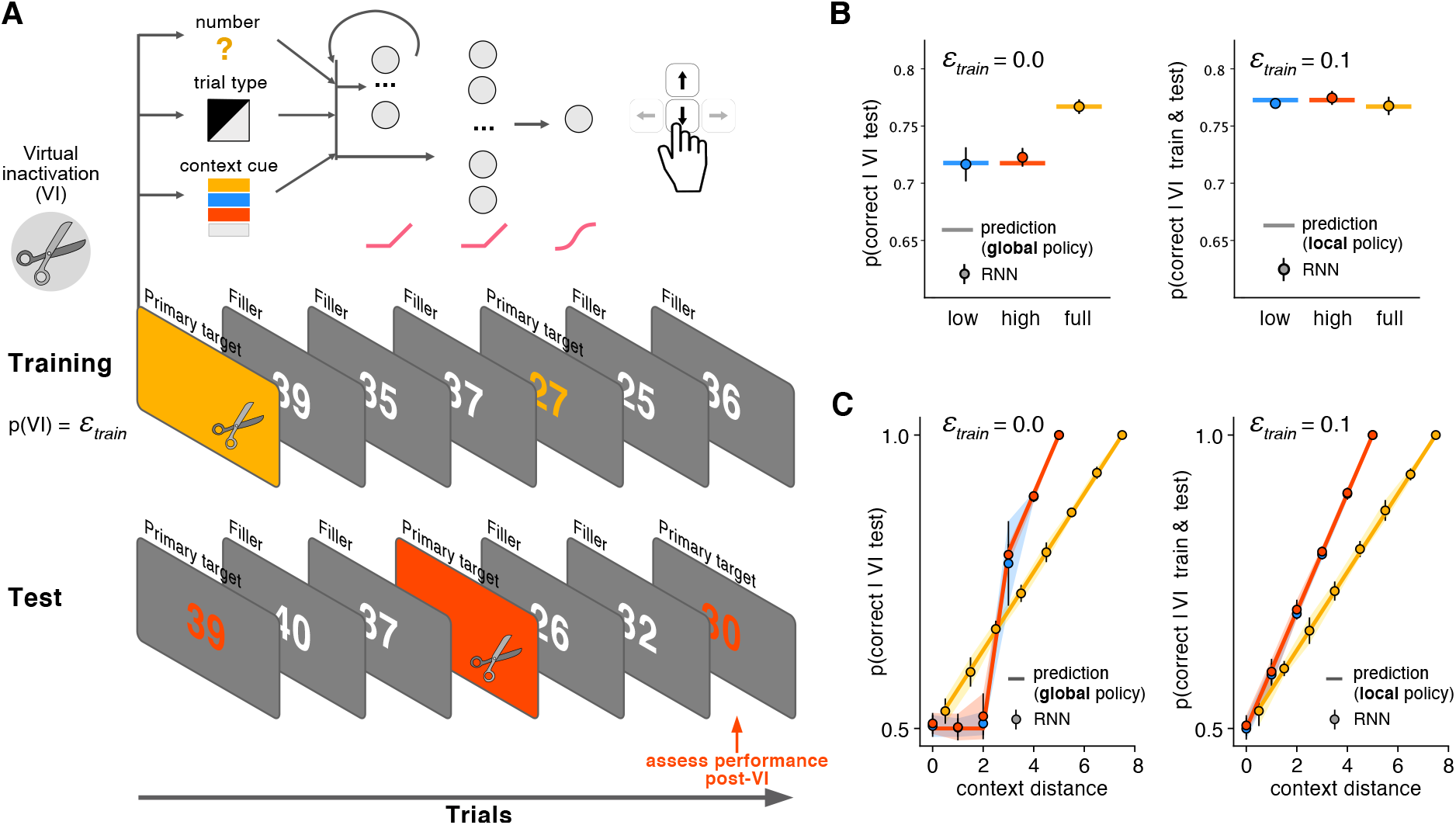
Recurrent neural network architecture, virtual inactivation and network behaviour. (A) Architecture of a recurrent neural network model trained to perform the same magnitude comparison task as humans. At each step the network received symbolic (one-hot) concatenated inputs indicating the number, trial type (filler or primary target), and context (zero for filler trials) of the current trial and indicated a binary “more” or “less” response with a single output node. The activation functions for each layer are shown in pink. Virtual inactivation (shown here with scissors) involved setting the number input on a given trial to zero, as if the number had been missed by the network. On each training trial, the network inputs corresponding to the primary target were virtual inactivated with probability ε_*train*_. At test, each primary target was virtually inactivated in turn and performance assessed on the primary target that followed. (B) Performance in each context predicted by an agent who optimally uses either the global (left, horizontal bars) or local (right, horizontal bars) context as a cue to magnitude comparison following a hidden (VI) target. RNNs (filled circles) without VI during training (left) and with VI during training (right) perform at exactly the levels predicted by the global and local agents respectively. (C) RNN accuracy (filled circles) following a VI shown as a function of local context distance for each context. RNN accuracy matches the predictions (coloured lines) of an agent using the global context (left) when the RNN was without VI in training (left). In contrast, the performance of RNNs with VI during training match predictions of an agent using the local context (right). Context colouring: low (blue), high (red), full (golden). Error bars in B and C show standard deviation across different random model initialisations and datasets. See also **Fig. S1**.

Assessing RNN performance on a subset of test trials for which the previous target input was virtually inactivated (ε_*test*_ > 0) offered information about how they were performing the task, both with and without VI during training. On these test trials, memory for the previous target is erased so that a network that ignores context will perform at chance. As a point of comparison, we computed the performance ceiling displayed by an agent who optimally uses either the local context (i.e. numbers within the current block) or global context (i.e. all numbers) as a lone cue to magnitude comparison. Networks that had undergone VI during training achieved near equivalent accuracy in low, high and full conditions, performing at precisely the level of an optimal agent using the local context (**Fig. 2b**). By contrast, networks that enjoyed perfect memory during training performed better on the full than high or low conditions, matching predicted performance levels for an ideal agent that used only the global context. In other words, whereas all networks learned to use context as a cue to respond, only when capacity was limited during training did the networks learn to exploit the range of numbers in the current context to maximise their accuracy; we confirmed this observation statistically by comparing the residuals of the fit to either model (both t-values > 16; p < 0.001 in both cases). In this and all subsequent statistical analyses, we treat the individual network (n = 10; each with their unique initialisation and stimulation sequence) as the unit of replication. In **Fig. 2c** we plot the network performance as a function of local context distance in each condition (low, high, full) for both networks trained with ε_*train*_ ∈ [0, 0.1] (see **Fig. S1** for ε_*train*_ ∈ [0.2, 0.3, 0.4]). When VI is applied, performance is a linear function of context distance in each condition, whereas without VI, performance remains at chance for context distances of ≤ 2 in the low and high conditions, because for these instances a comparison to the global average fails to offer the correct answer.

### Neural network state space

With these results in hand, we used representational similarity analysis (RSA) combined with multidimensional scaling (MDS) to visualise the embedding of numbers and contexts in the network neural state space. Focussing on fully trained networks, we computed correlation distance among hidden unit activations evoked by each number in each context and plotted the resulting neural states in three dimensions. In **Fig. 3a-b** we show the projection of each stimulus in each context (coloured dots) into a space spanned by the first three dimensions of the activations from the RNN hidden layer. As can be seen, the network learned to organise numbers according to their magnitude onto three parallel lines, one for each context. This occurs both with and without VI during training. However, consistent with the finding that local context is a more salient cue for responding when ε_*train*_ = 0.1, these context-specific number lines were more widely separated when VI was applied at training, as revealed by a statistical comparison of the Euclidean distance among their centroids (t_9_ > 25, p < 0.001).

**Fig 3.**
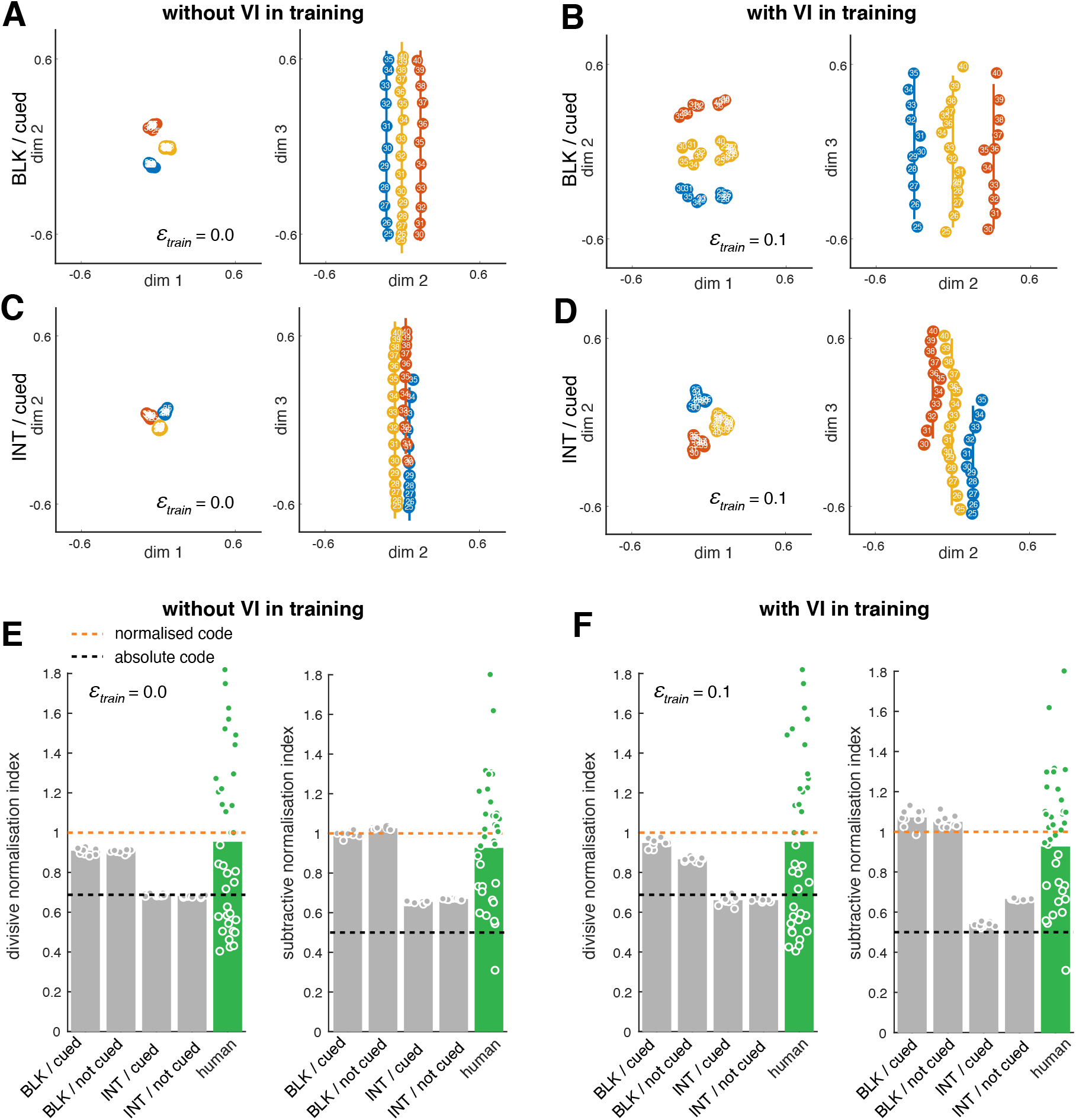
Neural network activations and normalisation metrics. Neural network models were trained either with or without virtual inactivation (VI), trained and tested with either temporally blocked (BLK) or interleaved (INT) contexts, and with contexts either cued explicitly in the input (cued) or not (not cued). (A-B) MDS visualisation of hidden unit activations in networks that were blocked during training and test and for which context was explicitly cued. (C-D) MDS activations from networks for which context was explicitly cued but for which contexts were temporally interleaved in training and test. Filled circles show the state space representation of stimuli in the low (blue), high (red) and full (golden) contexts; inset white numbers indicate the corresponding symbolic number input to the network. Coloured lines show the best fit lines model for quantifying the representational geometry. (E-F) Divisive and subtractive normalisation indices from fits to the representations which resulted from each network condition (grey), and from human neural recordings (green). Grey dots are fits to individual random model initialisations and datasets. Green dots are fits to individual human subjects (see below). Dashed horizontal lines indicate the divisive and subtractive normalisation indices expected under fully normalised (orange) and fully absolute (black) coding schemes.

Critically, however, it can also be seen that the neural representation of number in each context was compressed and centred so that the three lines span a common distance in one of the three dimensions of neural state space. This means that for the network, common positions on each line do not denote specific numbers, but rather encode abstract quantities corresponding to “more” or “less” within each context. Forming an abstract concept of “more” or “less” presumably facilitated the readout of signed context distance, permitting the network to solve the mapping problem, which may be particularly useful under the VI manipulation where capacity was limited. We note however, that this normalisation only occurred when the network received different ranges of numbers in three temporally distinct contexts (blocks), matching the task performed by humans. When inputs from different contexts were interleaved (but still signalled with a unique cue) the network failed to normalise, representing the signals in their native (absolute) frame of reference (**Fig. 3c-d**).

To quantify these observations, we fit a state space model to the RNN data representational dissimilarity matrices (RDMs, n = 10). The model was fit by varying the angle and length of three parallel number lines within a three-dimensional state space. Gradient descent was used to minimise their discrepancy with the neural state space data. This model, whose identifiability we verified using a parameter recovery approach (**Fig. S3**), allowed us to compute and compare the best-fitting line lengths in each context. Under a full (divisive) normalisation scheme, all three lines should have the same length (divisive normalisation [DN] index = 1), whereas without normalisation, the ratio of numbers in the low/high to full conditions should be ~0.69, reflecting their relative range (ranges of 16 vs. 11).

The relative line lengths are plotted on the leftmost bars **Fig. 3e** (without VI during training) and **Fig. 3f** (with VI). As can be seen, when magnitude ranges were separated into temporally distinct contexts (denoted BLK), the DN index approaches 1, irrespective of whether contextual cues are offered, and independent of whether VI was applied at training or not. When ranges were interleaved (INT), however, the DN index is ~0.69 or lower, indicative of an absolute code. In other words, without the benefit of blocked temporal context, the network does not use the context cue to distinguish among the different ranges. We also computed a subtractive normalisation (SN) index by comparing the offset in centroids among the high, low and full conditions along the principal magnitude coding axis. Again, blocking (but not interleaving) of contexts led to a full subtractive normalisation (SN ~1) whereby the centres of each line were brought into the same register, indicative of neural state space alignment. For completeness, we also fit models with the restriction that line lengths in each context should be equal (relative model) or that they should reflect the range of numbers in each context (absolute model). For (cued) BLK conditions, 10/10 RNNs were better fit by the relative model, whereas for (cued) INT conditions, 10/10 RNNs were better fit by the absolute model.

Normalisation of the number lines here should permit generalisation of magnitude information across contexts. To test for this in the RNNs, we trained a binary logistic regression model to classify stimuli as either “more” or “less” relative to the context mean. To assess generalisation, we used one context for training and a different context for evaluation (averaging over all pairs of contexts). We observed better generalisation for context-blocked networks than for context-interleaved network activations (context-blocked, 86% correct; context-interleaved, 69% correct; unpaired t-test, t_18_=13.11, p < 0.001; both with VI during training). These results were obtained using the naïve (high-dimensional) space given by the 200 network hidden units; the finding was replicated when we instead trained and tested classifiers using the dimensionality-reduced MDS activations (context-blocked, 84% correct; context-interleaved, 70% correct; unpaired t-test, t_18_ = 12.58, p < 0.001).

A parallel question is to what extent normalisation impacts the ability of networks to use the local context information. Specifically, why is it that networks trained with blocked context and no VI (**Fig. 3A**) and which formed normalised representations, did not use local context when we assessed their behaviour following VI at test (**Fig. 2**). We hypothesized that, although these networks form a neural geometry that would potentially allow them to use the local context, without VI during training they never learn an appropriate decoder that would permit this to be used for locally context-sensitive behaviour. We tested this by retraining the decoder alone (freezing all weights and biases except the final layer), and applying VI during retraining. Indeed, we found that after decoder retraining, networks exhibiting neural state space alignment (i.e. with blocked training and no VI) indeed learned to use the local context information (comparing residuals of the fit to local vs global models; paired t-test t19= −3.14, p=0.012) (**Fig. S4**).

These investigations of the RNN neural state space offered several insights about how transitive orderings (e.g. numerical magnitudes) may be represented in different contexts. First, networks embedded arbitrary inputs onto parallel number lines in a way that reflected their transitive ordering. When different ranges of numbers occurred in different temporal contexts, these number lines were normalised so that each embedding space stretched from “less” to “more” within that context. When VI was applied at training, the number lines spread out in dimensions orthogonal to the magnitude axis, and it was presumably this which facilitated the use of context-specific rather than context-general information to help solve the task. When we trained magnitude decoders, networks that formed normalised neural codes permitted cross-context generalisations of relative magnitude more readily than those which coded for the absolute numerical value of each stimulus.

### Human neural data

Finally, we turn to analysis of neural data recorded with scalp electroencephalography (EEG) whilst humans performed the numerical comparison task. Our main focus is the results of a multivariate analysis approach designed to interrogate the nature of the neural state space and its alignment across contexts. However, we also observed univariate signals with a right occipitoparietal focus that covaried positively with magnitude and with target distance in an early window (from 200-500 ms post-stimulus) and signals that covaried with local context distance over left posterior electrodes in a later window (500-800 ms post-stimulus). These effects, which survived correction for multiple comparisons using a familywise error test, are shown in **Fig. 4c**.

**Fig 4.**
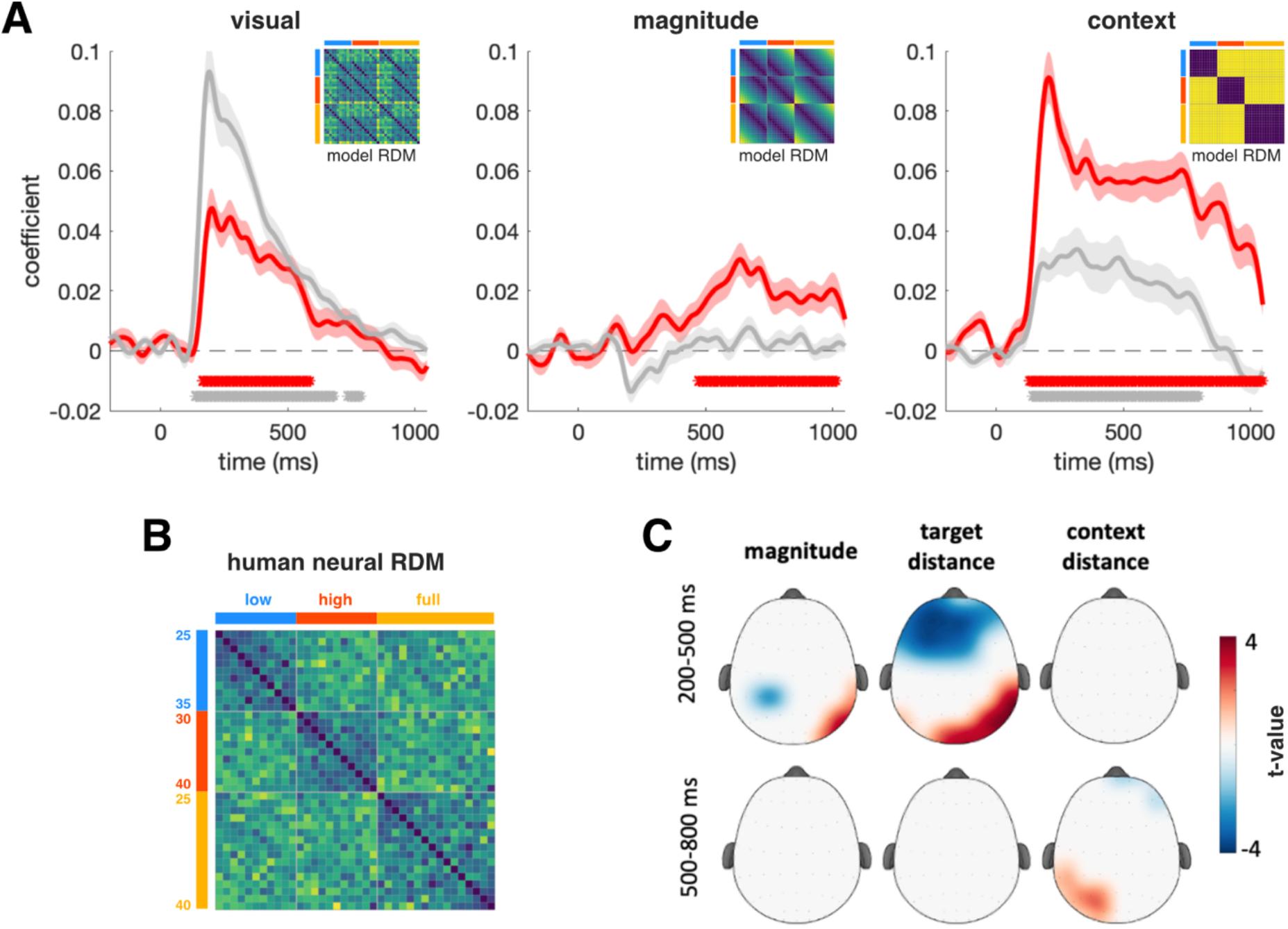
Human EEG encoding of context, magnitude and visual similarity. (A) Model representational dissimilarity matrices (shown inset) labelling the symbolic number within each context were regressed against the time course of neural activity within each trial (x-axis). The time course of regression coefficients is shown for both the filler trials (grey) and primary targets (red). Their cluster-corrected significance at each time point is indicated along the bottom of each figure. Coefficient time courses are shown resulting from regression against: an RDM indicating the pixel-wise visual similarity between inputs (left panel), an RDM indicating the numerical magnitude of each input (middle panel), and an RDM indicating the context (right panel). (B) A visualisation of the mean neural RDM across all participants. (C) Univariate effects of magnitude, target distance, and local context distance rendered onto the scalp. Warm colours indicate positive relationship; cold colours a negative relationship. Only effects that survived correction for multiple comparisons are shown. See also **Fig. S2**.

We constructed model RDMs on the basis of three variables: numerical magnitude, context, and the pixelwise similarity among inputs (Arabic digits). We then regressed these three model RDMs (standardised by z-scoring) against a data RDM obtained from scalp EEG patterns at each timepoint post-target; for a point of comparison, we also conducted the same analysis on the filler stimuli (**Fig. 4a**). As expected, early timepoints are dominated by visual similarity (left panel), for both targets (red lines) and fillers (grey lines). The visual signal is stronger for the filler numbers, which is likely to be due to the matching task participants performed on the fillers which required them to pay attention to the physical identify of adjacent filler stimuli. However, from about 600ms the EEG signals evoked by targets carried information about both context and magnitude; the context information was weaker, and the magnitude information was absent for the fillers. We also tried to decode responses to target stimuli (which were made with two fingers of the right hand) but were unable to do so at any timepoint across the epoch (all p-values > 0.2), making it unlikely that any partial correlation with response drives these or any subsequent results.

Next, we used MDS to visualise relations among the neural codes associated with each number in each context (**Fig. 5d, e**). Within each time bin, we fit a model that described the number relations with three parallel lines in neural state space. In **Fig. 5a**, we show how the slopes of those number lines vary over time using a sliding window approach and in **Fig. 5d**, we plot the first three dimensions of the neural state space in the early window (200-500 ms) and late window (500-800 ms). In the early period it can be seen that there is a reliable segregation of multivariate neural signals associated with each of the three contexts, with the neural distance between numbers within a context reliably smaller than those between contexts, compared to an appropriate shuffled null distribution (p < 0.001). Moreover, whilst there appears to be an emergence of three number lines (rightmost panel), the slopes of these lines did not fall within the top 5% of a null distribution constructed by shuffling the neural RDMs, and it can be seen that there is no alignment among the centroids of these lines.

**Fig 5.**
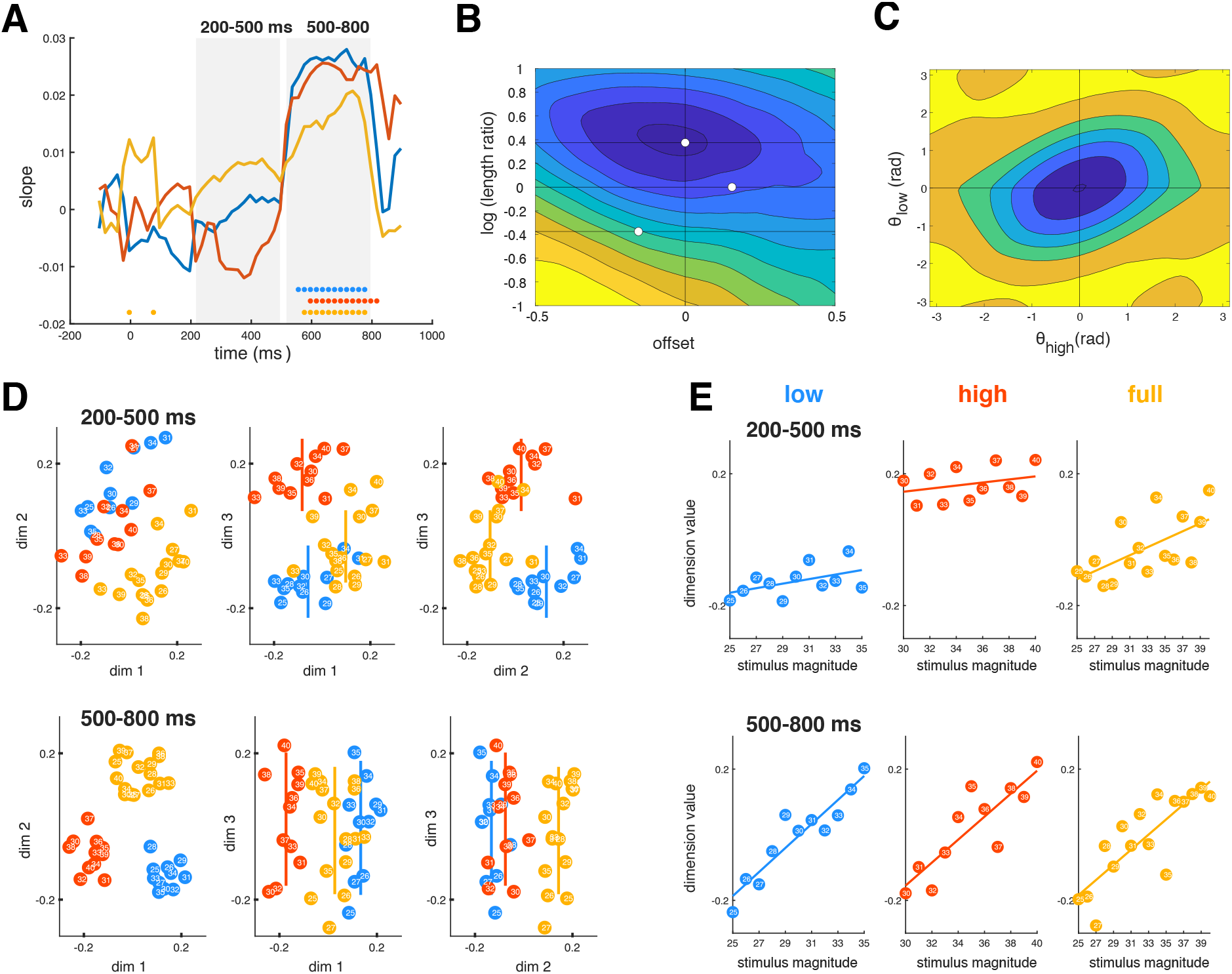
Human neural activity normalises the representation of magnitude. (A) Slopes of the relationship between number and dimension value calculated within a sliding temporal window for each context (blue, low; red, high; gold, full). Slopes were normalised by a shuffled control. Asterisks show reliable deviation from zero. (B) A contour map of the sum of squared residuals (averaged across the population) resulting from regressions of magnitude model RDMs that were constructed by parametrically varying the line centre offset (shown on x-axis) and relative length (y-axis) of each number line along the magnitude dimension. White dots indicate parameter combinations defining three specific models: relative coding (normalisation), absolute coding, and reverse coding (symmetric “anti-normalisation”). See also **Fig. S5**. (C) A contour map of the sum of squared residuals resulting from regressions of magnitude model RDMs constructed by varying the angle between the lines of 3 number lines of the same length. In both (B) and (C), blue pixels indicate better fits. (D) MDS visualisation of human neural activity corresponding to the primary targets shown in each context, mapped into a 3-dimensional space. Top panels, early window; bottom panel, late window. The neural codes associated with each number form equidistant clusters according to the context in which the number was presented (left panel). See also **Fig. S3**. (E) The position of each target along the third MDS dimension plotted against its numerical value. Each x-axis is scaled to indicate “small” and “large” numbers in each context. Plotted lines were fit to capture the slope of this relationship for each context and time-window. The slope of the best-fitting line is approximately equal in each context for the late time window only. Filled circles show the state space representation of stimuli in the low (blue), high (red) and full (golden) contexts; inset white numbers indicate the corresponding Arabic number shown on the trial. Coloured lines show the best fit lines model for quantifying the representational geometry.

However, as we move into the later time bin (lower panels) we can see that just as for the RNN, the numbers spread out along parallel lines in the third dimension, whereas the three contexts themselves were separated in the first two dimensions, lying approximately at the apices of an equilateral triangle (left panel). We also see a highly reliable effect of context as quantified by a permutation test (p < 0.001). The third dimension seemed to code for magnitude, with the lower numbers in each context exhibiting negative scores along this dimension and the higher numbers positive scores. To test this latter contention more formally, we assessed the rank correlations between the cardinalities and dimension value and found that they were much more positive than would be expected by chance, when compared to a shuffled control that randomly swapped the cardinality and number-dimension value assignments (Spearman’s rank correlation, p < 0.001 for all three contexts). We assessed the robustness of the association between cardinality and dimension-value by asking whether the relationship would hold if, in each context, the highest and lowest numbers were removed. When we excluded the most extreme (highest and lowest) numbers and repeated the Spearman’s tests, we found that the association continued to hold for all three contexts (all p-values < 0.03). We also plotted the magnitude of each number in each context against its value on the third dimension (**Fig. 5e**). Positive slopes were recovered in each case, and the slopes of the relationship between dimension scores and numerical magnitudes were more positive than expected by chance for all three contexts, as evaluated relative to a control that computed slopes from shuffled number-dimension value assignments (all p-values < 0.001).

Next, to quantify the geometry of these representations, we created parametric families of RDMs and regressed them against the neural data for each participant individually, mapping how well systematic variation in the representations of numbers and contexts explained the data. We began by fitting a simplified model that assumed that number relations were described by lines of equal length (full normalisation), and used this model to assess whether number lines were more parallel than expected by chance in neural state space. As the lines are elongated along the third MDS dimension only, we collapse the data across the first two dimensions to consider the simplified case of 2D rotation of the low and high context lines (with angles denoted *θ*_*low*_ and *θ*_*high*_) with respect to the full. We then predicted the neural data as a weighted sum of the “visual” and “context” model RDMs shown in Fig 4a, as well as a “magnitude” RDM constructed from these two angles. We parametrically varied the angles that were used to construct magnitude model RDM and obtained the best-fitting solution for each participant. We observed that the best-fitting line rotations were more concentrated around 0° than would be expected by chance (Rayleigh test, p < 0.006; *θ*_*low*_: *R* = 9.66, *p* < 0.015; *θ*_*high*_: *R* = 11.37, *p* < 0.006), and in **Fig. 5c** we show the fit deviance across all participants has a clear minimum at the origin, i.e. (*θ*_*low*_, *θ*_*high*_) ≈ (0,0). In other words, the best fit overall is a model in which the three lines (low, high, full) are on average almost exactly parallel.

The abscissa in **Fig. 5e** is scaled such that relatively “small” and “large” numbers are spaced equally in the three contexts, and under this scaling, it can be seen that the slope of the best-fitting line is approximately equal in each context. This implies that the number lines are normalised within a context to span the same distance in neural state space, in a way that (in theory) would allow for generalisation of abstract information – “more” vs. “less” – between contexts. We tested this effect statistically in a number of different ways. Firstly, we fit number lines to the MDS plots for individual participants (as we did for the RNNs above), using the resulting estimates to compute the indices for divisive normalisation (DN; line length relativisation) and subtractive normalisation (SN; relative centering of the lines). The average DN index for humans was 0.95 ± 0.41, which was statistically indistinguishable from 1 (the value expected for a normalised code; p = 0.76) but greater than the relative length ratio 11/16 (expected under an absolute code; t_37_ = 3.96, p < 0.001). Similarly, the value for SN was 0.93 ± 0.38, which was indistinguishable from 1 (expected for a normalised code; p = 0.88) but greater than 0.5 (expected under an absolute code; t_37_ = 6.84, p < 0.001). These results are depicted via the green bars in **Fig. 3e-f**.

Secondly, building on the approach described above, we fit model RDMs that parametrically varied both the relative length and offset between each line and attempted to predict the neural data for each individual participant. To verify that this any apparent relativisation in the number lines was not spuriously related to normalisation implicit in MDS, we conducted a parameter recovery exercise, which showed that our model could recover the precise form of the mental number lines under the assumptions of both a relative and an absolute code (**Fig. S3**). In **Fig. 5b** we plot the average sum of squared residuals as a function of these parameters to show that the global minimum does indeed fall almost exactly on the point predicted by the relative (normalisation) model. When we used Bayesian model comparison for group studies (Stephan et al., 2009) to compare the relative, absolute and reverse models, we find strong evidence for the relative model, with exceedance probabilities indistinguishable from 1 and the majority of participants (80.92% expected frequency) explained by the relative model over the other two models. Moreover when we compared models across the full space of line length ratios, we found strong evidence in favour of a model with log(*k*_*low,high*_) ~0.4, which corresponds more or less exactly to that expected under a fully relative coding scheme (**Fig. S5**). This suggests that as for RNNs, the neural number lines are normalised in a way that facilitates neural state space alignment. For completeness, we also regressed the neural network models against the human EEG data, finding that a model with VI during training, cueing and blocking fit best (**Fig. S3**).

We note in passing that this neural state space alignment implies that humans represent numbers in each context with a relative code rather than an absolute code. This allows us to make a further prediction about behaviour: that the relative (but not absolute) difference between successive numbers should be the best predictor of decision accuracy and response times. We thus computed the rectified difference in magnitude between two successive primary targets (coded as absolute or relative values) and regressed these models competitively to find that the relative distance model positively predicted participant accuracy (t-test on group-level *β*_*rel*_, t_37_ = 2.99, p < 0.004), and negatively predicted reaction times (t_37_ = −3.26, p < 0.002). However, the absolute distance model did not predict either (*β*_*abs*_, t-values < 1).

Finally, returning to the neural data, we tested directly whether the neural codes in humans supported the generalisation of magnitude across contexts, conducting our analysis on the average RDM across participants (**Fig. 4c**) and projecting the data into either the three-dimensional MDS space (**Fig. 5d**) or the highest dimensional space extractable from the RDM (38-dimensional). We then trained binary logistic regression models in the same way as for the neural networks and compared generalisation performance to classifiers trained and tested on label-shuffled data. We found that these neural signals indeed supported significant generalisation of abstract magnitude when trained and tested on the dimensionality-reduced data (3-dimensional data, p < 0.001), and with a more marginal significance, the full-dimensionality data (38-dimensions, p < 0.03 one-tailed). However, generalisation was not significant when trained and tested on the raw electrode data at the individual participant-level (60-dimensional raw electrode data, one-sided p=0.323), presumably because the unprocessed signals recoded at the scalp are more noisy.

## Discussion

We studied the neural representation of number and context in humans and neural networks performing a sequential magnitude comparison task. Consistent with previous reports, we found that human decisions were guided by local contextual signals as well as memory for the previous item. This implies that humans maintain an estimate of the local average number within a block to help guide responding when memory is weak. The context-related decision information (“context distance”) is encoded in univariate neural signals over occipito-parietal electrodes, albeit in a rather late window beginning at approximately 500 ms post-stimulus. These findings are broadly consistent with previous reports that where imperative information is weak or absent, judgments are biased towards the central tendency of the stimulation history (de Gardelle & Summerfield, 2011; Holllingworth, 1910; Jazayeri & Shadlen, 2010) and that (in the rodent) this contextual information is coded in parietal circuits (Akrami et al., 2018). Recurrent neural networks trained to perform the task do not naturally use local contextual information when their memory is fully intact. However, when their memory is artificially rendered fallible during training, the networks also learn to capitalise on local contextual information to make judgments of relative magnitude in a way that closely resembles the human participants.

Our major question was the nature of the neural representation of magnitude and context in humans and neural networks. We studied this by reducing the dimensionality of neural signals recorded at multiple scalp electrodes in humans, and hidden unit activations read out from neural networks trained to perform the task, visualising and quantifying the neural state spaces in which each number and context was embedded. The most striking finding is the highly conserved way that humans and neural networks represent magnitude and context in this task. In both model systems, magnitudes are projected onto parallel neural “number lines” whereby stimuli with greater magnitude difference elicited more dissimilar neural signals in each context. Moreover, in both systems these number lines are normalised so that they span a common distance within neural state space, meaning that the ends of each line correspond to “more” and “less” – a relational quantity, rather than a specific numerical value.

A low-dimensional code for symbolic number has been reported before in M/EEG signals (Luyckx et al., 2019; Spitzer et al., 2017; Teichmann et al., 2018). This neural code must be abstracted away from the physical properties of the inputs, because physical similarity among Arabic digits is not determined by their relative cardinality. Here, we used two-digit numbers, and so there was some unavoidable correlation between magnitude and physical similarity due to decade boundaries (e.g. 29 and 30), but we still observed a multivariate code for number even after regressing out pixelwise similarity among digits (**Fig. 4b**), and no such code was observed for comparable filler objects, so we think it is unlikely that putative magnitude effects are driven by visual appearance. Similarly, because we were unable to decode the response made by participants to target stimuli (with two fingers of the right hand), we think it is unlikely that putative magnitude effects are related to motor signals.

However, rather than a generic code for number, we observed that both humans and neural networks learned to additionally segregate information by context, so that rather than a single number line, we observed three parallel neural number lines. In humans, this pattern did not occur instantly but emerged gradually over the course of the epoch. Early in the epoch (e.g. from 200 ms) neural signals distinguished context itself, an effect that may have been driven in part by the different font colour in which targets occurred across blocks. During this early window we observed nascent number lines in distinct parts of state space, but without parallel arrangement. From 500ms, however, numbers were arranged into context-specific parallel number lines.

We argue that this neural geometry has at least two desirable properties. Firstly, it allows number information in each context to potentially be kept separate. This is useful because (as discussed above) when memory is imperfect the local context provides helpful information about how to respond; if all numbers were projected onto a single line, this contextual information would be unavailable. Indeed, when we examined the state spaces of neural networks that had not experienced virtual inactivation during training, and that did not use local context distance as a cue, the three parallel number lines were much closer together. In fact, when training was interleaved so that time could not be used as an additional cue for context, they lay virtually on top of one another (**Fig. 3**).

Secondly, the fact that number lines are parallel in neural state space facilitates generalisation between physically similar stimuli that share a common magnitude. Irrespective of whether individual neurons exhibit specialised coding or exhibit mixed selectivity, this neural geometry ensures that a linear decoder trained to classify stimuli according to their magnitude in one context could be successfully applied to read out magnitudes in another, thus permitting cross-context abstraction (Bernardi et al., 2020). Indeed, a long tradition emphasises that humans generalise naturally between space, time and number, by using a magnitude representation that is shared across different input modalities (Hubbard et al., 2005; Walsh, 2003). This theory also successfully predicts the existence of neurons that code for magnitude with a shared code across input domains; such cells have been identified in the parietal cortex of macaque monkeys experiencing trains of auditory or visual pulses (Nieder, 2012). In humans, the parietal cortex is a hub for numerical cognition (Piazza & Izard, 2009) and a site where overlapping representations of “distance” signalled by spatial, temporal or social comparisons are observed in fMRI signals (Parkinson et al., 2014). Similarly, when humans learned to rank images of animals according to their probability of paying out a reward, shared multivariate patterns in EEG signals come to code for the number and value, as if there were a corresponding neural signal for higher numbers and higher event probabilities (Luyckx et al., 2019). Together, these findings imply a general principle whereby neural state space alignment permits generalisation across contexts (Bernardi et al., 2020).

However, generalisation of relational information between contexts can be hampered by what we call the “mapping problem”: the need to represent stimulus geometry on a common scale. For example, our task requires participants to estimate whether a number is reflective of “more” or “less” in each context in case memory is imperfect. In recurrent neural networks, we can see that this is achieved as the neural number lines are both subtractively and divisively normalised so that they span a common distance along one axis of neural state space, meaning that this axis runs from “more” to “less” rather than simply indexing numerical value. This normalisation should improve generalisation: indeed, when we tested this, we found that in both humans and neural networks that state space alignment (for example those that enjoyed blocked training) it was possible to generalise the notion of “more” vs. “less” by training a decoder in one context and testing it in another. However, generalisation was poorer in those neural networks whose geometry was not indicative of neural state space alignment. Of note, no normalisation is explicitly engineered into our networks; rather, they autonomously learn to encode numbers in this context-normalised fashion because doing so enhances performance.

In humans, we observed evidence that supported the same normalisation process: the neural number line in the “full” condition appeared to be compressed to a similar length in neural state space as that for the “low” and “high” conditions (divisive normalisation), and the centroids of the numbers were aligned along one dimension (subtractive normalisation). It has long been known that biological brains are prone to normalise incoming sensory signals, with neural activity typically expressed relative to the local average response (Carandini & Heeger, 2012). Explanations for this ubiquitous motif often focus on the need to make efficient use of computational resources (Louie et al., 2013). For example, divisive normalisation helps make efficient use of the dynamic range of a neuron or population, and subtractive normalisation can “explain away” redundant information in an input signal (Rao & Ballard, 1999). This theory predicts that numbers should be coded within a common range in neural state space, but does not dictate that they should be parallel, as observed here, or allow generalisation between contexts as our decoding analysis (discussed above) implies. Here, we suggest a role for normalisation (complementary to any role in efficient coding) in aligning neural codes to facilitate generalisation across contexts. Other studies offer hints of a comparable process. For example, when monkeys reproduced either long or short temporal intervals (contexts), the underlying neural dynamics in the medial prefrontal cortex were found to ‘stretch’ or scale in time according to context (Wang et al., 2018).

These suggestive findings notwithstanding, we acknowledge that our data do not allow us to impute a direct causal link between neural state space alignment and the deployment of the generalisable concept of “more” or “less” for generalisation – or, in the current study, to facilitate performance on the target matching task. We observed that neural networks trained and tested under blocked conditions with VI (who exhibited neural state space alignment) use the local context when information about the past target is unavailable. However, the picture is nuanced by the data from the networks trained *without* VI – who formed closely-overlapping but normalised neural number lines, but still failed to use these to display local context-sensitive behaviour. This shows that while neural state space alignment may be necessary for cross-context generalisation, it is not sufficient for the networks to use the local context. This is presumably because whilst this neural geometry in theory permits the read out of an abstracted code for “more” and “less”, it can only do so in the presence of an appropriate decoder. Consistent with this explanation, when we retrained the decoder of context-blocked networks with no VI, and applied VI during this retraining, the networks learned to display context-sensitive behaviour (**Fig. S4**).

Our study leaves a number of other questions unanswered. Firstly, we focus here on a very simple form of relational abstraction – the transitive ordering implied by numerical magnitudes. It remains to be seen whether the results described here generalise to more complex relational structures. Secondly, because of the limited spatial resolution of EEG, and the use of multivariate methods that relied upon electrodes from across the scalp, we are unable to say much about the neural locus of our effects. On the basis of the univariate findings (which highlight occipito-parietal electrodes) and past work (Akrami et al., 2018), we think that the parietal cortex is a likely candidate for representing magnitude in parallel, context-specific lines. However, we cannot say this with confidence on the basis of the current data. Finally, we note that normalisation was not ubiquitous across the human cohort. Within the late time window, a relative model fit better in a majority of participants but was not ubiquitous. It would be interesting to conduct more targeted tests to ask whether participants that normalise more effectively also generalise more readily. These caveats aside, however, we believe that these findings offer insights into the neural coding and generalisation of the concept of magnitude, a basic form of abstraction for humans (Walsh, 2003).

## Author Contributions

F.L., C.T. and C.S. conceived human experiments. F.L. and C.T. collected human behavioural and EEG data. H.S. and C.S. conceived neural network modelling. H.S. implemented neural network modelling. H.S., F.L., S.N. and C.S. conceived and implemented analyses. H.S., F.L. and C.S. drafted the paper. H.S, F.L, S.N and C.S. edited and revised the paper.

## Acknowledgements

This work was funded by the European Research Council (grant REP-725937 to C.S.) and received funding from the European Union’s Horizon 2020 Framework Programme for Research and Innovation under Specific Grant Agreement 785907 (Human Brain Project SGA2, to C.S.)

## STAR Methods

### RESOURCE AVAILABILITY

#### Lead Contact

Further information and requests should be directed to and will be fulfilled by the Lead Contact, Hannah Sheahan

#### Materials Availability

This study did not generate new unique materials.

#### Data and Code Availability

Data is available upon request. Code for the RNN simulations and analyses is available on github at https://github.com/hannahsheahan/context_magnitude, and the code for the human data analyses is available on github at https://github.com/summerfieldlab/Number_lines.

### EXPERIMENTAL MODEL AND SUBJECT DETAILS

Thirty-nine human participants were recruited for the experiment through the recruitment system at the Department of Experimental Psychology at the University of Oxford. One participant was omitted from the analyses due to technical difficulties with the EEG equipment. All analyses were performed on the remaining 38 participants (15 female, 23 male, age = 27.11 ± SD 6.13). Participants were required to have normal or corrected-to-normal vision, with no history of neurological or psychiatric illness. Participants were compensated for their time at a rate of £10 per hour, with an additional maximum bonus of £5 determined by their performance in the task in which they achieved lower accuracy (see below). The reward contingency was added to ensure participants would pay equal attention to both tasks. Informed consent was given before the start of the experiment. The study was approved by the Medical Science Inter-Divisional Research Ethics Committee (R49432/RE001).

### METHOD DETAILS

#### Experimental procedure

Stimuli were created and presented using the Psychophysics Toolbox-3 (Brainard, 1997; Kleiner et al., 2007) for Matlab (MathWorks) and additional custom scripts. The tasks were presented on a 20-inch screen with a resolution of 1600 x 900, at a refresh rate of 60 Hz and on a grey background. Viewing distance was fixed at approximately 62 cm. The ‘up’ and ‘down’ arrow keys on a standard QWERTY keyboard were used as response keys for the numerical comparison task and the space bar for the number matching task.

The experiment was a dual-task rapid serial visual presentation paradigm, involving a numerical comparison task and a number matching task (see **Fig. 1**). In each block (n = 24), participants viewed a sequence of 120 two-digit numbers. Numbers relevant for the numerical comparison task (“targets”) were presented in coloured font, while those relevant for the number matching task were presented in white (“fillers”). On presentation of a target number, participants were asked to compare its magnitude to that of the previous target number in the stream, responding ‘up’ when it was greater and ‘down’ when it was smaller using their right hand. Every target was followed by 2 - 4 fillers. Participants were asked to press space bar with their left hand when a filler number was identical in magnitude to the previous target. This secondary task was imposed to keep participants focussed on all numbers.

The event sequence was as follows. Each block started with the presentation of a central fixation cross (1000 ms) followed by two placeholder hashtag signs (500 ms). Each stimulus (number) was presented for 500 ms with a fixed ISI of 1000 ms after a target and a variable ISI (750-1250 ms) after a filler. Participants were free to respond from stimulus onset until 200 ms before the onset of the next stimulus (this avoided feedback signals overlapping with the presentation of the subsequent stimulus). If a response key was pressed, participants received auditory feedback for 150 ms. Correct responses were followed by a high-pitch tone and all errors resulted in a low-pitch tone. If participants failed to respond within the appropriate time window, a buzz sound was presented for 150 ms. At the end of each block, participants were informed about their percentage accuracy on both tasks.

One each block, targets (for the numerical comparison task) were drawn from a specific range. In the **Low** range context, numbers were uniformly drawn between 25 and 35, in the **High** range context between 30 and 40 and in the **Full** range context between 25 and 40. The white filler numbers always spanned the entire range between 25 and 40 irrespective of block. The experiment consisted of 8 blocks of each context. Each context was associated with a unique colour for the targets (blue: RGB = [86, 180, 233]; orange: RGB = [230, 159, 0]; purple: RGB = [230, 120, 220]) and the colour-to-context mapping was randomised between participants. The block order was randomised in order to reduce temporal similarity between numbers from the same context. Each block contained 29 targets (excluding the first coloured number of the sequence), with a probability of 12% that at least one of the subsequent white numbers matched the previous target. To prevent potential task switching costs, the white filler number immediately after a primary target never required a response, but participants were not made aware of this feature. At the start of the experiments, participants first completed 3 practice blocks of 144 trials each, one for each of the three contexts. These blocks did not count towards their final performance bonus.

#### EEG acquisition

The EEG signal was recorded using 61 Ag/AgCl sintered surface electrodes (EasyCap, Herrsching, German), a NeuroScan SynAmps RT amplifier, and Curry 7 software (Compumedics NeuroScan, Charlotte, NC). Electrodes were placed according to the extended international 10-20 system, with the right mastoid as recording reference and channel AFz as ground. Additional bipolar electrooculography (EOG) was recorded, with two electrodes placed on either temple for recording horizontal EOG and two electrodes above and below the right eye for vertical EOG. All data was recorded at 1 kHz and low pass filtered online at 200 Hz. All impedances were kept below 10-15 kΩ during the experiment.

#### EEG pre-processing

The data were pre-processed using functions from the EEGLAB toolbox (Delorme & Makeig, 2004) for Matlab and custom scripts. First the data were down-sampled to 250 Hz, low-pass filtered at 40 Hz and then high- pass filtered at 0.5 Hz. The continuous recording was visually screened for excessively noisy channels and these were interpolated by the weighted average of the surrounding electrodes. The data was then offline re-referenced to average reference. Epochs were extracted from 250 ms before to 1000 ms after stimulus onset. Epochs were baselined relative to the full pre-fixation time window. Epochs containing atypical noise (such as muscle activity) were rejected after visual inspection. We then performed Independent Component Analysis (ICA) and removed components related to eye blink activity and other artefacts (manually selected for each participant).

#### RNN architecture

A simple recurrent neural network model was built to perform the same primary task as human participants. The network was trained to transform symbolic inputs **x** on each trial t using a recurrent layer ***h***^*(1)*^, followed by a fully connected feed-forward layer ***h***^*(2)*^, which projected to a single output node y. Rectified linear (ReLU) activation functions *f(u)* were applied to each of the two hidden layers and a sigmoid activation function *σ(u)* was applied to the output layer to constrain the response 0≤y≤1. Thus the network took the form:

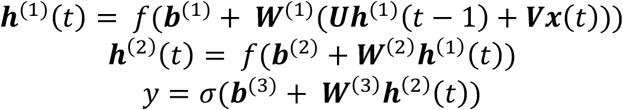

where ***U*** and ***V*** are binary matrices used for concatenating the current input ***x**(t)* and the recurrent hidden activations from the previous trial *h*^*(1)*^*(t-1)*. Vectors ***b***^(i)^ are the biases on layer I, and the two hidden layers ***h***^*(1)*^ and ***h***^*(2)*^ were 220 and 200 units wide respectively. Weights and biases on all layers were initialised with random samples from a uniform distribution spanning ±1/sqrt(n), where n is the number of nodes in the upstream layer. The hidden state on the first trial in the first block of each training and test set was initialized to zero, and subsequent blocks were initialised with the final hidden state of the previous block. Neural networks were built and trained in PyTorch (www.pytorch.org)

#### Training the network

Data was generated to simulate the trial sequence presented to the human participants, and so followed the same generation procedure. As in the human study, each block was 120 trials in length, containing 30 primary targets drawn from numbers corresponding to a single context, while filler trials always spanned the full range. The order of blocks was pseudorandom and the number of blocks in each context was balanced. However, unlike the human participants, the network had no a priori knowledge of numerical magnitude, and so many more blocks of trials (2880 blocks) were generated for training each network than the 24 blocks used per human participant. The RNN was trained only to perform the primary task, and responses on the filler trials were ignored.

On each trial, the input vector **x** consisted of a one-hot representation of the current number, as well as a node indicating whether the current trial was a primary target or a filler trial, and an (optional, see Context Manipulations) one-hot coded vector reflecting the current context. The context cue was included to simulate the colouring of the primary targets by context range in the human experiment. As filler trials were always white in the human experiment, the context cue inputs to the network were always zeroed on filler trials. The primary target numbers input to the RNN for each context were sampled from the same distributions as in the human experiment but were represented by one-hot codes spanning the ranges 1-16 (full range), 1-11 (low range), and 6-16 (high range) to save on input nodes.

The network was trained to perform the primary task (numerical comparison) and during training received feedback on primary target trials according to a binary cross-entropy cost function (but no feedback was given on the first primary target in each block). Errors were backpropagated through time at the end of each block of trials and the weights updated with stochastic gradient descent. Network outputs >0.5 were interpreted as analogous to pressing the ‘up’ arrow key in the human experiment, meaning that the current number was thought to be greater than the previous primary target while responses <=0.5 were interpreted as presses of the ‘down’ arrow key. Training hyperparameters were fixed for all networks at 10 epochs, a learning rate 0.0001, and momentum of 0.9. Network parameters were frozen at the end of training, prior to test.

Each network was trained 10 times, with different random initialisations and random datasets. Each dataset consisted of a training set (2880 blocks), and a test set for assessing the network activations (480 blocks).

#### Context manipulations

To isolate factors that could lead to context-separation and normalisation in the RNN activations, we factorially controlled the provision of explicit context cues to the network (analogous to colour in the human experiment), and the blocking of trials in time by numerical context. We trained four groups of networks to fully cross these two factors. In each group of networks, context cue inputs on primary trials either reflected the numerical range of the block, or these inputs were kept constant across all blocks. Additionally, primary targets were either drawn from a single numerical range within a block, as in the human experiment, or primary targets within each block were drawn from the global distribution of primary target numbers, which spanned all three numerical ranges. Therefore, while the blocking of numerical range changed between groups of networks, the total number of times each number occurred as a primary target across the dataset remained the same for all groups. For each network, the same manipulations were made across training and test sets.

#### Virtual inactivation

Primary target trials were virtually inactivated (VI; zeroing the inputs that communicated number) randomly with probability ε during training, and errors backpropagated. Groups of networks were trained with one of four different VI probabilities, ε = {**0.0, 0.1, 0.2, 0.3, 0.4**}. Context was explicitly cued in the input for all networks with VI during training. To evaluate whether the trained networks learned to use numerical context when solving the primary task, we used VI on a single primary target in a test set with temporally blocked contexts and assessed performance on the subsequent primary target. In effect, this forced the network to ‘guess’ whether the numerosity of the current input was likely to be greater or less than a ‘forgotten’ previous primary target. Network weights were frozen prior to any VI assessments on the test set. This process was repeated for all primary targets in the test set and post-VI performance was taken as the mean percentage of trials that the network answered correctly, evaluated on the primary target trial after the VI.

#### RNN activations

Network activations were evaluated on the test set, and for each condition the test set had the same structure as its associated training set (with the exception of VI which was not applied while network activations were measured). Inputs were passed to the network as in training, and the activation of units in the final hidden layer **h**^(2)^ was recorded and these activations averaged across all presentations of the same number, context and trial type (primary, or filler) in the test set. The geometry of these activations was assessed separately for each trained model (**Fig. 3e-f**) and the activations shown in **Fig. 3a-d** were computed by taking the mean activations across all 10 models trained under each condition before MDS was performed.

#### RNN decoder retraining

We assessed whether, given the pressure of imperfect memory, the normalised representations formed by networks which were originally trained without VI, but with temporal blocking (**Fig. 3A**), could be repurposed for context-dependent magnitude judgements. We froze the weights and biases of all trained layers except the final layer, reinitialised the weights of this layer only and retrained the network for 30 epochs, applying VI to alternate primary targets. In other words, we retrained the decoder for the representations shown in **Fig 3A** with temporally blocked contexts and VI so that the network could benefit from local context use. After retraining, we repeated the analysis performed on our original networks in which we used a paired t-test to compare the residuals between the network’s behaviour under VI at test and theoretical performance limits under local and global context use. We repeated this retraining procedure for networks originally trained under interleaved contexts and no VI, retraining under temporally blocked contexts with VI. We then compared the residuals between the behaviour of both sets of networks and a theoretical local context policy with an unpaired t-test. The results of these analyses are shown in **Fig S4**.

### QUANTIFICATION AND STATISTICAL ANALYSIS

#### Behavioural analysis

We modelled participant accuracy on primary targets using a logistic regression model with predictors for the disparity between current and past target number (*target distance*), the distance between the target number and the median number in the current block (*local context distance*) and the distance to the overall median number (across all blocks; *global context distance*). To test whether imperfect memory was what drives the use of context, we also compared the influence of context on accuracy when there were either 2, 3 or 4 intervening filler stimuli. We included an additional term in the previous regression to encode the interaction between number of fillers and the local context.

In order to directly compare behavioural models that are based on the “absolute” and “relative” coding schemes, we performed a complementary analysis in which we standardised numbers in two ways: either across the entire experiment (preserving the differences in mean and standard deviation between contexts) or within each context (so that numbers within a context had the same mean and deviation). We then used these variables to compute the absolute difference in magnitude between two successive targets and asked whether it was possible to exploit the residual variance to pose the question of whether numerical distance in absolute or relative space best explained RT or accuracy.

We entered these predictors into a regression for each participant individually, using logistic regression for accuracy, and log-transforming RT to correct for the rightward skew:

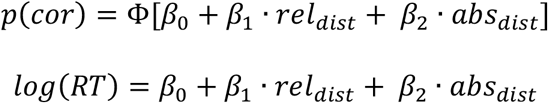

We then conducted group-level statistics using one-sample t-tests on *β*_1_ and *β*_2_ in each case.

#### EEG: Univariate analyses

A regression-based approach allowed us to disentangle the influence of various variables on the univariate signal during the numerical comparison task. Before running the regression, we used Principal Component Analysis (PCA) on the pre-processed data of each participant to reduce noise in the signal. All epochs with primary targets were stacked over all electrodes and trials, creating a feature matrix of (trials*electrodes) by time points. The first 43 principal components (PC) were retained, which on average explained 90% of the data. The EEG data was then reconstructed by multiplying the PC’s with the estimated loadings and the reconstructed data was subsequently used as the dependent variable in our linear regression model.

The regression model contained 4 regressors of interest: (1) numerical magnitude of the current primary target, (2) the absolute difference of the current primary target to the previous target, (3) absolute difference between the *current* target and the mean of the current context, and (4) absolute difference between the *previous* target and the mean of the current context. We included two nuisance regressors to exclude alternative explanations of the univariate results: (5) visual similarity of the current primary target with the previous target and (6) reaction time (RT) on the current trial. Visual similarity was estimated as the correlation distance between black-and-white pixel images of two numbers as they were presented on screen. Trials with no response or RTs beyond 2.5 SD of the inverse RT were excluded from analysis. All regressors were z-scored before estimating the beta coefficients for each time point and electrode. For a control analysis, the same regression model was repeated replacing the regressors coding for distance to the context mean with the distance to the global mean for current and previous target.

#### EEG: Representational Similarity Analysis (RSA)

We constructed neural Representational Dissimilarity Matrices (RDMs) from the EEG data at each time point. First, data was z-scored over all trials per electrode and time point. Condition-specific neural signals were estimated at each time point and electrode using a regression model with dummy codes for every number in each context (in total 38 predictors: 25-35 for Low, 30-40 for High and 25-40 for Full), where the beta coefficients reflected the average deflection in EEG signal per condition at each electrode. The residuals of the regression were used to estimate the covariance matrix, which was subsequently used to increase the signal-to-noise ratio (SNR) by noise normalizing (whitening) the averaged EEG data through multiplication with the negative half inverse covariance matrix (Σ^−0.5^) at each time point (Walther et al., 2016). Finally, we calculated the Pearson correlation distance between each condition, resulting in a 38 x 38 RDM at each time point. To exclude the possibility that more observations in certain contexts were biasing the dissimilarity measures, we constructed RDMs iteratively with subsampled data that equated the frequency of observations in each cell of the RDM. We repeated the RDM construction 100 times, randomly selecting N observations per cell at each iteration, with N determined by the minimum number of observations in any condition for a particular participant. The final neural RDM (*nRDM*) then consisted of the averaged RDM over all 100 iterations.

We constructed 3 candidate model RDMs that represented different potential structures in the neural signal: context, magnitude and visual similarity. The context model RDM assumed complete similarity (0) between numbers coming from the same range context and complete dissimilarity (1) for numbers coming from different ranges. The magnitude model RDM encoded the absolute difference between each number in each context. The visual similarity RDM was calculated using correlation distance between the vectorized black-and-white pixels of the numbers as they appeared on screen. The regression had the following form:

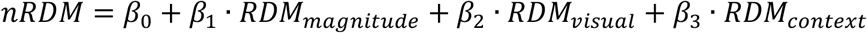

All model RDMs were z-scored before entering in the regression to assess their relative contribution. Beta coefficients were estimated at each time point and for each participant separately and the resulting beta series were smoothed over time for visualisation. Statistics are reported at the group level. Significant clusters were identified using cluster-corrected non-parametric permutation tests (Maris & Oostenveld, 2007).

For a supplementary control analysis, two additional subject-specific models were added to the regression: a colour model RDM and a reaction time (RT) model RDM. For the model RDM representing the colour space, RGB values were first transformed into CIE 1976 L*a*b* space to more accurately reflect human perception of colour differences. The dissimilarity between colours was then indexed through ΔE∗, a measure of the colour difference between two colours in CIELAB space. The RT RDM was constructed by calculating the average RT to a number independent of the preceding number. RTs were cut-off based on the 2.5 standard deviation from the inverse RT to control for outliers before averaging.

#### Representational geometry with RSA

We performed regressions that attempted to predict the neural data RDM (*nRDM*_*i*_) for each human participant *i*. The predictors were a set of z-scored RDMs (similar in form to those inset in **Fig. 4a** and described above) that collectively encode the similarity between the neural signal evoked by each number in each context. The regression has the following form:

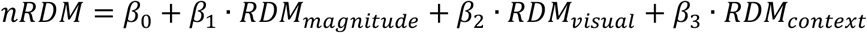

To quantify the representational geometry of the neural number lines, we vary *RDM*_*magnitude*_ parametrically in a way that allows us to estimate the best-fitting number line length for each participant. Specifically, we model the numbers in each context *c* as lying in an ordered and evenly spaced fashion on a line ranging from −*k*_*c*_ + *m*_*c*_ to *k*_*c*_ + *m*_*c*_ on the third dimension of our simulated neural space. In other words, *k*_*c*_ determines the line length and *m*_*c*_ determines the offset of the line centre from the origin in each context.

For the full context, we assume *k*_*full*_ = 16/11; for the high and low context, we assume *k*_*low*_ = *k*_*high*_ but otherwise allow the parameter to vary freely. Because *k* determines the ratio of line length of the low/high to full conditions, it is natural to vary it logarithmically. Thus, models where *k*_*low*_ = *k*_*high*_ = 1 [or log(*k*_*c*_) = 0] assume that numbers are represented on an absolute scale (with no normalisation). By contrast, values of *k*_*low*_/*k*_*high*_ that are greater than 1 are indicative of normalisation (with 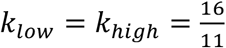 indicating full normalisation); those that are less than one indicate the converse, a sort of (perhaps counterintuitive) reverse normalisation. Under this scheme, thus, if best-fitting values of *k*_*low*_/*k*_*high*_ are reliably greater than 1 at the group level then this indicates evidence for normalisation.

Simultaneously, we allow *m*_*c*_ to vary in the range [−0.5, 0.5], assuming that *m*_*high*_ = −*m*_*low*_ and *m*_*full*_ = 0. This means that the lines in the low and high context are offset in opposite directions from the origin with the line for the full context centred intermediate between the two. Thus, if *m*_*high*_ = −*m*_*low*_ = 0 then the lines are all centred at the same point. This is what is predicted by the normalisation model. Note that if *k*_*full*_ = 16/11 and 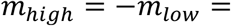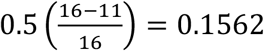 then each point on each line is equated for magnitude. This is predicted by a model proposing that neural codes for number are not normalised within context.

Systematically varying the values of *m*_*c*_ and *k*_*c*_, for each model variant we construct *RDM*_*magnitude*_ as the Euclidean distance between each number and every other number in each context. Substituting this RDM in the equation above, for each participant, we estimated coefficients exhaustively (using grid search) for values of *m*_*c*_ and *k*_*c*_. The sum of squared residuals (i.e. the model deviance) of the resulting regressions are shown below as a contour map, averaged over the population, with blue pixels indicating lower values (better fit) in **Fig. 5b**. We then used Bayesian model comparison for group studies (Stephan et al., 2009) to compare the evidence for relative, absolute and reverse models. We also compared the full space of models defined by log(*k*_*c*_) by exhaustively testing points along the line of offset=0 (**Fig. S5**).

#### Assessment of parallelism with RSA

We develop further the regression methods described above to assess the angles between the neural number lines. To deal with the potentially high-dimensional search space involved in fitting freely oriented lines, we make some simplifying assumptions. Firstly, building on the results above, we assume that the numbers are spaced regularly and in order on lines of equal length (full normalisation). Secondly, because the lines are elongated along the third principal component only, we collapse over the first two dimensions, reducing our problem to one of 2D rotation (rather than 3D). We can also fix the orientation of the number line for the full condition (aligned to the 3rd dimension) and rotate the lines from the low and high conditions freely, because there exist identities in the solution that allows all 3 lines to rotate. These simplifications allow us to define two parameters *θ*_*low*_ and *θ*_*high*_, which pertain to the angle of the low and high number lines with respect to the full. We then use these rotations to define *RDM*_*magnitude*_ and use an identical regression-based approach to identify the best fit to the neural RDMs at the per-individual participant level:

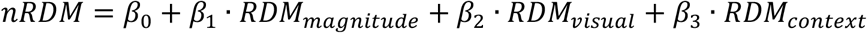

Because we have only 2 parameters, we can visualise the deviance (sum of squared residuals) on a surface (**Fig. 4**). We subsequently use a Rayleigh test to show that based on individual participant fits, the best-fitting rotations are more concentrated around 0° than would be expected by chance.

#### Model fitting to low-dimensional data

Neural state spaces were visualised by reducing the similarity data (RDMs) to three dimensions using multidimensional scaling with metric stress (equivalent to plotting the first three principal components of the data). Next, we used a model fitting exercise, complementary to the method described above, to characterise how the neural representation of number and context is organised in this low-dimensional space.

To start with, we assumed that each context was characterised by a neural representation of numbers lying equally spaced on a line, that the centroid of the [x,y,z] coordinates for the numbers in each context was the midpoint of this line, and that the three lines (one for each context) were parallel. We fit a model with 6 degrees of freedom: parameters 1-3 were the lengths of the each of the three lines, and parameters 4-6 were the angles of the (parallel) lines in dimensions [x,y,z]. The model was fit using gradient descent to minimise the discrepancy between simulated number positions and observed number positions in the low-dimensional neural state space. We used parameter recovery to verify that this model could recover ground truth line lengths and orientations in a simulated space (**Fig. S3**). We used this approach both for human data (see **Fig. 4**) and RDMs constructed from RNN hidden unit activations (see below and **Fig. 3**). We used the same approach on the group mean similarity data (average RDM) and individual human/network RDMs. For each subject (n = 38 humans; n = 10 RNNs), we fit three variants of the model: one in which the line lengths were constrained to be the same (relative model); one where the line length in the full condition was constrained to be 16/11 times longer for the full condition than low or high (reflecting the larger range; absolute model); and one where no constraints were imposed (free model). The residuals (loss) of the absolute and relative models were compared using frequentist statistics.

The line length parameters estimated from the free model were used to compute indices of divisive normalisation as follows:

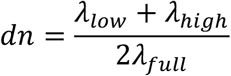

where *λ*_*i*_ is the estimated length parameter for condition *i*. The index *dn* will thus be 1 for perfect normalisation, i.e. when the line lengths in full and low/high conditions are equal and will approach 11/16 if the line length is reflective of the absolute range of numbers in each context. The subtractive normalisation index was calculated as follows:

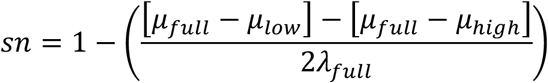

where *μ*_*i*_ is the centre of dimension *d* for condition *i*. This index captures the relative offset of the centroid of the low/high conditions from the full condition, normalised by the line length in the full condition. This index will approach 1 if there is full normalisation (i.e. if there is no offset) and 0.5 for full offset. We ensured that any meaningful offset occurred in dimension ***d* = 3** by rotating the lines (using the best fitting parameters 4-6) so that any elongation on a magnitude axis, if present, would occur in the dimension 3 (we also visualised neural state spaces after this rotation had been applied).

#### Assessment of number ordering

The model fitting described above in which we regressed the magnitude RDM against the neural data captures the ordering of the stimuli across all three number lines collectively. We also performed additional analyses to test whether the cardinality of each number was encoded along the third neural dimension in each individual line. We computed the slope of the relationship between the number cardinalities and the dimension value in the group mean neural data, and for each context compared this to an empirical null distribution consisting of 10,000 random permutations that swapped the number-dimension value assignments. We also computed Spearman’s rank correlation as a measure of the association between variables because of its robustness and reduced sensitivity to outliers and compared these to 10,000 random permutations that swapped number-dimension assignments.

As a complementary analysis, we assessed whether the association between cardinality and dimension value would hold if, in each context, the highest and lowest numbers were removed. We tested this in the same fashion as described above, although it should be noted that the number of possible permutations for comparing the relationship in the data to is reduced.

#### Relating behaviour to neural geometry

We modelled both reaction time and accuracy behaviour with relative and absolute models, and obtained estimates of goodness of fit as the squared error of the regression models including only a single term (either relative or absolute):

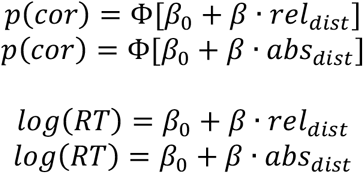

This yielded 4 estimates of goodness of fit, which we correlated with the model evidence from the absolute and relative models of neural geometry at the individual participant level (modelling of neural geometry is explained in the Methods section *Multidimensional Scaling and Model Fitting*). We used Spearman’s rank correlation to minimise the influence of outliers. The results showed that there was variance in RT which related to the neural geometry. Specifically, the fit of the relative behavioural model predicted the fit of the relative neural model, and the fit of the absolute behavioural model predicted the fit of the absolute neural model (all Spearman’s *ρ* > 0.5, all p < 0.001). No such relationship was observed for accuracy. This strong association validates our overall neural modelling approach.

However, the *difference* in fit of the relative and absolute behavioural models did not predict the *difference* in fit of the neural models. In other words, this was a general, but not a specific association – it is not the case, as far as we can tell, that those participants whose neural data were relatively better fit by a relative model were also those whose reaction times were best fit by a relative model. This is likely to be because the different predictions of the relative and absolute model are quite subtle, and it is hard to distinguish them at the single-subject level.

#### Calculations of normative performance

To evaluate the usage of local context information by the RNN, we compared the post-VI RNN performance to two benchmark levels. These benchmarks were calculated as the best overall performance achievable for an agent following each of two different policies when presented with a primary target (and not the previous primary target). These were πlocal: a policy that used the local context information when responding. Under policy πlocal the agent guesses the current primary target *x*_*A*_ to be greater than the previous (inactivated, forgotten) primary target *x*_*B*_ if the current target is greater than the median of the current range of primary targets, assuming the range of primary targets was learned during training. We also evaluated performance under πglobal: a policy that used the global context when responding. Under policy πglobal the agent guesses that the current primary target *x*_*A*_ was greater than (inactivated, forgotten) *x*_*B*_ if *x*_*A*_ is greater than the median of all primary targets across all contexts (which was similarly learned during training). That is

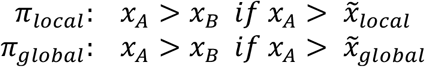

The probability that an agent following policy π responds correctly on a random trial is given by

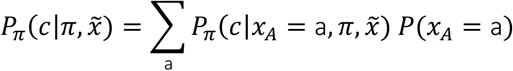

For either the local or global case, applying one of the above policies gives

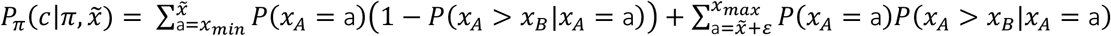

Under a context-blocked training and test regime, the distribution of *x*_*A*_ is uniform across the local context range of primary targets (full, low or high), and the distribution of *x*_*B*_ is uniform across the same range with the exception that x_A_ ≠ x_B_. If NA is the number of possible outcomes of x_A_, and NB the number of possible outcomes of x_B_, then

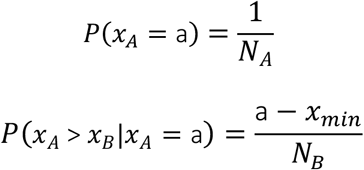

The performance values presented in **Fig. 2 & S1** were then found by substituting values for x_A_ for each context and policy.

#### Generalisation tests

To assess whether the representations formed by the neural networks and observed in the EEG data permitted magnitude to generalise across contexts, for each context we trained a binary logistic regression model to classify stimuli as larger or smaller than the context mean. In the networks the input to these models were the activations produced by the final hidden layer in response to stimuli in each context (the dimensionality-reduced data is displayed in **Fig. 3**). Models were trained on activations corresponding to a single context, and we tested generalisation performance on the other two contexts. We repeated this exercise for all context folds in each RNN, training and testing the big/small classifier on either the dimensionality-reduced MDS activations (3-dimensional), or the full set of activations across all 200 hidden units (200-dimensional). For each trained network (10 per condition), we calculated the mean generalisation performance across all context combinations and used an unpaired t-test to compare the scores between networks that had temporally blocked contexts to those for which contexts were temporally interleaved.

We repeated this analysis on the human EEG data. We first tested this on a subject-by-subject level, attempting to cross classify stimuli in one context after training on the data from other contexts. We repeated the analysis on the group average data (obtained from averaging RDMs across participants), and quantified our results by constructing an empirical null distribution using 10,000 classifiers trained on the same EEG data, but with stimuli labels shuffled within each context (we cannot compare to interleaved as we did for neural networks, because we do not have the data for the humans). We repeated this analysis using both the low-dimensional (3-dimensional) and high-dimensional (38-dimensional) group average data.

#### Model recovery

To verify that our MDS pipeline yielded accurate results, we used a parameter recovery approach. We created synthetic data in which each number was characterised by a simulated neural code that resembled that predicted by the “absolute” model. In other words, each stimulus (number) was defined by a position in a coordinate space whose value on one dimension (dim 3) was the same for all numbers of equivalent absolute magnitude (e.g. numbers 30 in low context and 30 in high context share the same coordinate value on that dimension) and whose coordinate on a different dimension (dim 2) is determined by its context. We added noise to these synthetic inputs and used them to create an RDM, exactly as we did for human neural data. Following exactly the processing pipeline adopted for neural data, we then asked whether MDS could recreate the pattern of synthetic data we began with.

We specified the centre of the lines in the low, high and full contexts at [0, -c, m], [0 0 -m]; [0 c 0] respectively, where c signals the offset along dim 2 for each context, and m signals any offset from the centre of the full line (along dim 3) in the low and high conditions. We first attempted to recover the absolute model by setting c = 0.1 and m = 0.15. We define the spread of the number codes in the 3rd dimension as [0.2 0.2 0.2*k] in low, high, full (where k = 16/11, i.e. the line in the full is longer by a factor of 16:11). We add a small amount of noise to each neural pattern, and find that when we visualized the MDS, it indeed recapitulated the pattern in the synthetic data (**Fig. S3**).

#### Correspondence between RNN and human neural RDMs

We took RDMs from all 8 neural network conditions, generated by varying the following factors: (i) no VI vs. VI during training, (ii) blocked vs. interleaved, and (iii) cued vs. not cued. We regressed these against the whole-brain data RDM at each post-stimulus timepoint in turn. Within the late period (500 – 800 ms post-stimulus) we saw the largest beta coefficients (slopes) for the model with VI during training, cueing, and blocked training (**Fig. S3C**).

To test for statistical significance, we calculated the residuals of the fit to each condition within the 500-800ms period and entered them into a 2 x 2 x 2 ANOVA. This yielded a three-way (VI x cue x blocking) interaction F_1,37_ = 6.3, p < 0.02, qualifying the idea that VI during training, cueing and blocking were all required for a good fit between human and model data.

## References

Akrami, A., Kopec, C. D., Diamond, M. E., & Brody, C. D. (2018). Posterior parietal cortex represents sensory history and mediates its effects on behaviour. Nature, 554, 368–372.

Baram, A. B., Muller, T. H., Nili, H., Garvert, M., & Behrens, T. E. J. (2019). Entorhinal and ventromedial prefrontal cortices abstract and generalise the structure of reinforcement learning problems. BiorXiv Preprint. https://www.biorxiv.org/content/10.1101/827253v1

Behrens, T. E. J., Muller, T. H., Whittington, J. C. R., Mark, S., Baram, A. B., Stachenfeld, K. L., & Kurth-Nelson, Z. (2018). What Is a Cognitive Map? Organizing Knowledge for Flexible Behavior. Neuron, 100, 490–509.

Bellmund, J. L. S., Gardenfors, P., Moser, E. I., & Doeller, C. F. (2018). Navigating cognition: Spatial codes for human thinking. Science, 362.

Bernardi, S., Benna, M. K., Rigotti, M., Munuera, J., Fusi, S., & Salzman, C. D. (2020). The Geometry of Abstraction in the Hippocampus and Prefrontal Cortex. Cell, 183, 954–967.

Brainard, D. H. (1997). The psychophysics toolbox. Spatial Vision, 10, 433–436.

Carandini, M., & Heeger, D. J. (2012). Normalization as a canonical neural computation. Nat Rev Neurosci, 13, 51–62.

Collins, A. G., & Frank, M. J. (2013). Cognitive control over learning: Creating, clustering, and generalizing task-set structure. Psychol Rev, 120, 190–229.

de Gardelle, V., & Summerfield, C. (2011). Robust averaging during perceptual judgment. Proc Natl Acad Sci U S A, 108, 13341–13346.

Delorme, A., & Makeig, S. (2004). EEGLAB: An open source toolbox for analysis of single-trial EEG dynamics including independent component analysis. Journal of Neuroscience Methods, 134, 9–21.

Doumas, L. A., Hummel, J. E., & Sandhofer, C. M. (2008). A theory of the discovery and predication of relational concepts. Psychol Rev, 115, 1–43.

Ernst, M. O., & Banks, M. S. (2002). Humans integrate visual and haptic information in a statistically optimal fashion. Nature, 415, 429–433.

Fitzgerald, J. K., Freedman, D. J., Fanini, A., Bennur, S., Gold, J. I., & Assad, J. A. (2013). Biased associative representations in parietal cortex. Neuron, 77, 180–191.

Ganguli, S., Bisley, J. W., Roitman, J. D., Shadlen, M. N., Goldberg, M. E., & Miller, K. D. (2008). One-dimensional dynamics of attention and decision making in LIP. Neuron, 58, 15–25.

Gentner, D. (2010). Bootstrapping the mind: Analogical processes and symbol systems. Cogn Sci, 34, 752–775.

Holllingworth, H. L. (1910). The central tendency of judgement. The Journal of Philosophy, Psychology and Scientific Methods, 7, 461–469.

Hubbard, E. M., Piazza, M., Pinel, P., & Dehaene, S. (2005). Interactions between number and space in parietal cortex. Nat Rev Neurosci, 6, 435–448.

Jazayeri, M., & Shadlen, M. N. (2010). Temporal context calibrates interval timing. Nat Neurosci, 13, 1020–1026.

Kleiner, M., Brainard, D., & Pelli, D. (2007). “What’s new in Psychtoolbox-3?” Perception, 36.

Kording, K. (2007). Decision theory: What “should” the nervous system do? Science, 318, 606–610.

Kording, K. P., & Wolpert, D. M. (2004). Bayesian integration in sensorimotor learning. Nature, 427, 244–247.

Lake, B. M., Ullman, T. D., Tenenbaum, J. B., & Gershman, S. J. (2017). Building machines that learn and think like people. Behav Brain Sci, 40, e253.

Lake, Brenden M., Salakhutdinov, R., & Tenenbaum, J. B. (2015). Human-level concept learning through probabilistic program induction. Science, 350, 1332–1338.

Louie, K., Khaw, M. W., & Glimcher, P. W. (2013). Normalization is a general neural mechanism for context-dependent decision making. Proc Natl Acad Sci U S A.

Luyckx, F., Nili, H., Spitzer, B., & Summerfield, C. (2019). Neural structure mapping in human probabilistic reward learning. Elife, 8.

Maris, E., & Oostenveld, R. (2007). Nonparametric statistical testing of EEG- and MEG-data. J Neurosci Methods, 164, 177–190.

Murphy, G. L. (2002). The Big Book of Concepts. MIT Press.

Nieder, A. (2012). Supramodal numerosity selectivity of neurons in primate prefrontal and posterior parietal cortices. Proc Natl Acad Sci U S A, 109, 11860–11865.

Parkinson, C., Liu, S., & Wheatley, T. (2014). A common cortical metric for spatial, temporal, and social distance. J Neurosci, 34, 1979–1987.

Piazza, M., & Izard, V. (2009). How humans count: Numerosity and the parietal cortex. Neuroscientist, 15, 261–273.

Rao, R. P., & Ballard, D. H. (1999). Predictive coding in the visual cortex: A functional interpretation of some extra-classical receptive-field effects. Nat Neurosci, 2, 79–87.

Remington, E. D., Egger, S. W., Narain, D., Wang, J., & Jazayeri, M. (2018). A Dynamical Systems Perspective on Flexible Motor Timing. Trends Cogn Sci, 22, 938–952.

Spitzer, B., Waschke, L., & Summerfield, C. (2017). Selective overweighting of larger magnitudes during noisy numerical comparison. Nat Hum Behav, 1.

Stephan, K. E., Penny, W. D., Daunizeau, J., Moran, R. J., & Friston, K. J. (2009). Bayesian model selection for group studies. Neuroimage, 46, 1004–1017.

Summerfield, C., Luyckx, F., & Sheahan, H. (2019). Structure learning and the posterior parietal cortex. Prog Neurobiol, 101717.

Teichmann, L., Grootswagers, T., Carlson, T., & Rich, A. N. (2018). Decoding Digits and Dice with Magnetoencephalography: Evidence for a Shared Representation of Magnitude. J Cogn Neurosci, 30, 999–1010.

Tenenbaum, J. B., Kemp, C., Griffiths, T. L., & Goodman, N. D. (2011). How to grow a mind: Statistics, structure, and abstraction. Science, 331, 1279–1285.

Tervo, D. G. R., Tenenbaum, J. B., & Gershman, S. J. (2016). Toward the neural implementation of structure learning. Curr Opin Neurobiol, 37, 99–105.

Walsh, V. (2003). A theory of magnitude: Common cortical metrics of time, space and quantity. Trends Cogn Sci, 7, 483–488.

Walther, A., Nili, H., Ejaz, N., Alink, A., Kriegeskorte, N., & Diedrichsen, J. (2016). Reliability of dissimilarity measures for multi-voxel pattern analysis. NeuroImage, 137, 188–200.

Wang, J., Narain, D., Hosseini, E. A., & Jazayeri, M. (2018). Flexible timing by temporal scaling of cortical responses. Nat Neurosci, 21, 102–110.

