## Supplemental Figures for "Neural state space alignment for magnitude generalisation in humans and recurrent networks"

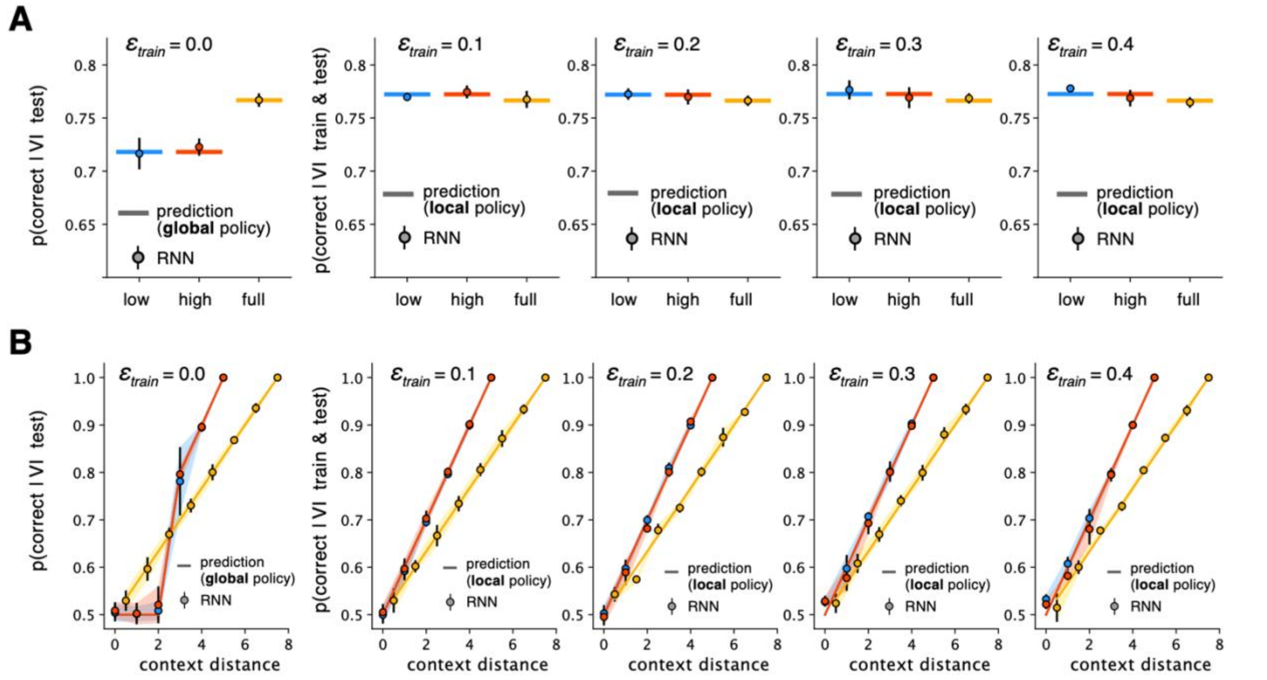


**Fig S1. Neural network behaviour for different rates of virtual inactivation during training (related to Fig. 2).** As in Figure 2, neural network performance at test assessed on the primary target immediately following virtual inactivation, shown here for different frequencies of VI during training, $\varepsilon_{train}$. (A) Performance in each context predicted by an agent (horizontal bars) who optimally uses the global (left most panel) or local (all other panels) context for magnitude comparison following a VI target. RNNs (filled circles) without VI during training (left most panel) and with VI during training but at different frequencies (all other panels). (B) RNN accuracy (filled circles) following a VI and predictions of local and global agents (coloured lines), plotted as a function of local context distance for each context. While networks without VI during training match the predictions of an agent using the global context (leftmost top and bottom panels), all tested networks with $\varepsilon_{train} \geq 0.1$ match predictions of an agent using the local context as a cue in magnitude comparison. Context colouring: low (blue), high (red), full (golden). Errorbars show standard deviation across different random model initialisations and datasets. N = 10 models for each VI frequency level $\varepsilon_{train} < 0.2$, N = 4 for each level $0.2 \leq \varepsilon_{train} \leq 0.4$.

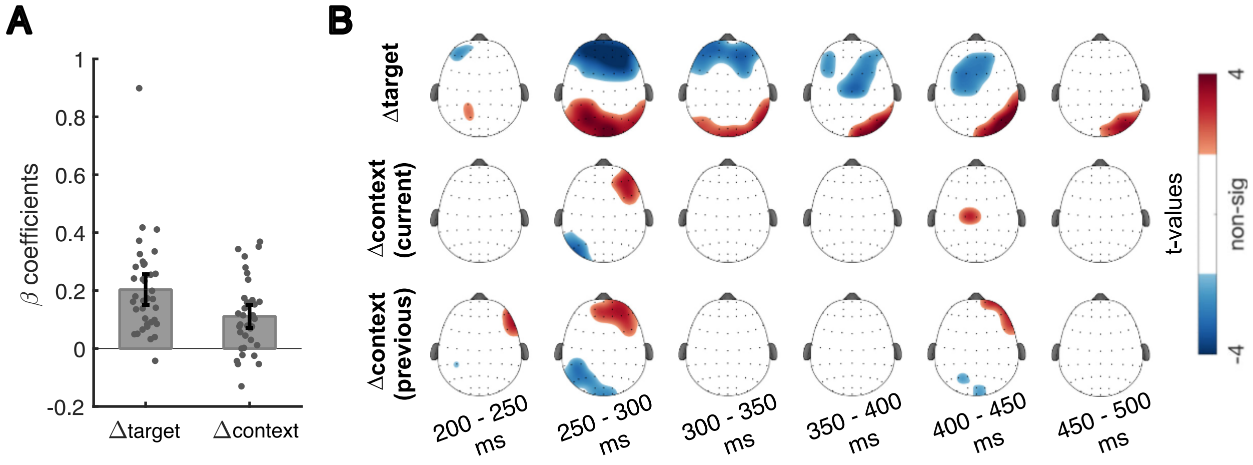


**Fig S2. Univariate analysis of human neural activations (related to Fig. 4)**. (A) Regression coefficients from regressing target and context distance predictors onto single-trial EEG data. (B) The beta series mapped over time across the scalp show target distance (Δ target, unsigned difference between the two target numbers) was positively encoded over posterior parietal electrodes from ~250-400ms post-stimulus. A consistent representation of local context distance (Δ context current) between the currently observed target and the local context median was observed over right prefrontal electrodes. A similar signal was observed for the previous target (Δ context previous), as if participants encoded whether each item was "more" or "less" independent of its specific identity.

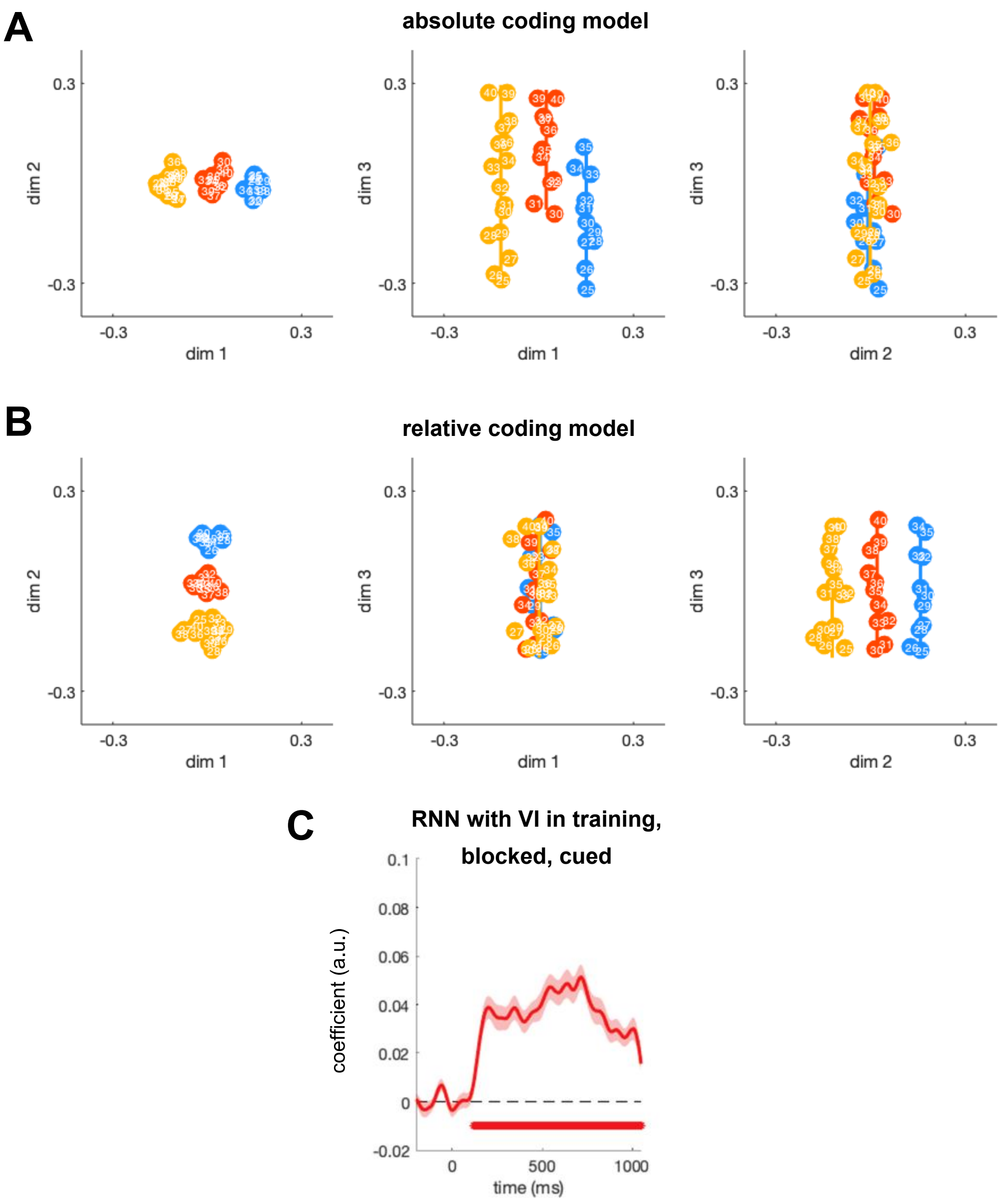


**Fig S3. Model recovery and correspondence between human and RNN RDMs (related to Fig. 5).** Model RDMs with parametrically varied line lengths and context separation were fit to both human and RNN activations to quantify their geometry. When we generated fake data RDMs under both an absolute coding scheme (A) and a relative coding scheme (B), and added a small amount of noise to each pattern, we recovered MDS plots that captured the appropriate lengths and offsets of the lines in each context. (C) Coefficients of a regression between the whole-brain human data RDM at each timepoint and the RNN model with VI, contexts cued, and blocked training. The red bar shows timepoints for which there was a significant linear relationship. Out of all 8 neural network conditions ((i) no VI vs. VI during training, (ii) blocked vs. interleaved, and (iii) cued vs. not cued), this is the RNN model for which we observed the largest beta coefficients (slopes) during the 500-800ms period.

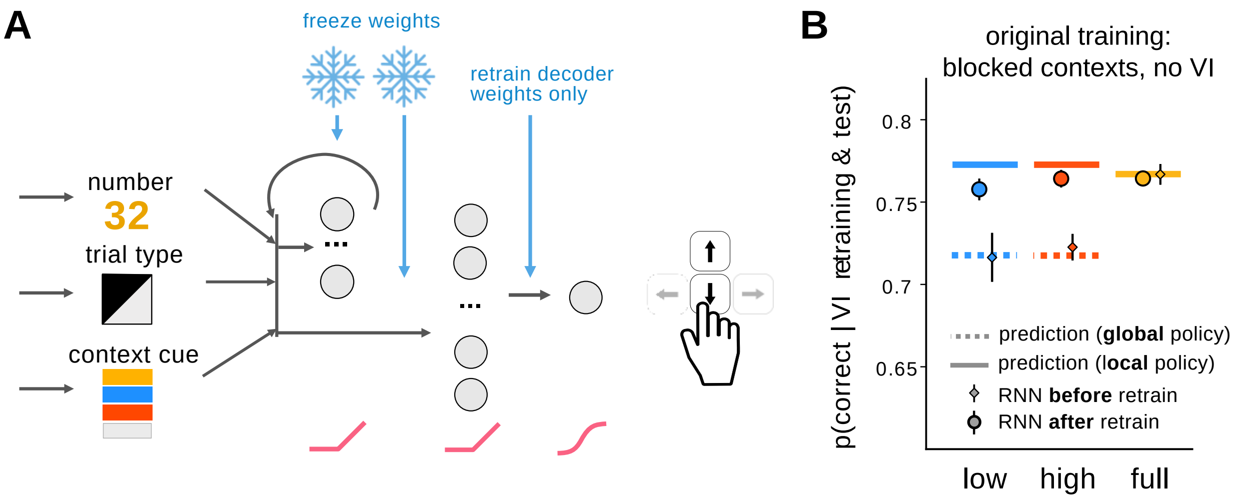


**Figure S4. Local context use for network decoders retrained with VI.** (A) All weights and biases were frozen in the RNNs except for the final hidden layer, which were reinitialized and retrained under temporally blocked contexts with VI. (B) Local context use was assessed as before: at test, each primary target was virtually inactivated in turn and performance assessed on the primary target that followed. Local context use was assessed on a temporally context-blocked test set before (small diamonds) and after (circles) retraining. RNNs (filled circles) originally trained with blocked contexts and no VI, but which had the decoder weights retrained under blocked contexts with VI, learned to use local context. The theoretical policy which best fits the retrained network data (local policy) is shown with a solid horizontal line for each context, and the alternative global policy is shown with a dashed horizontal line for reference.

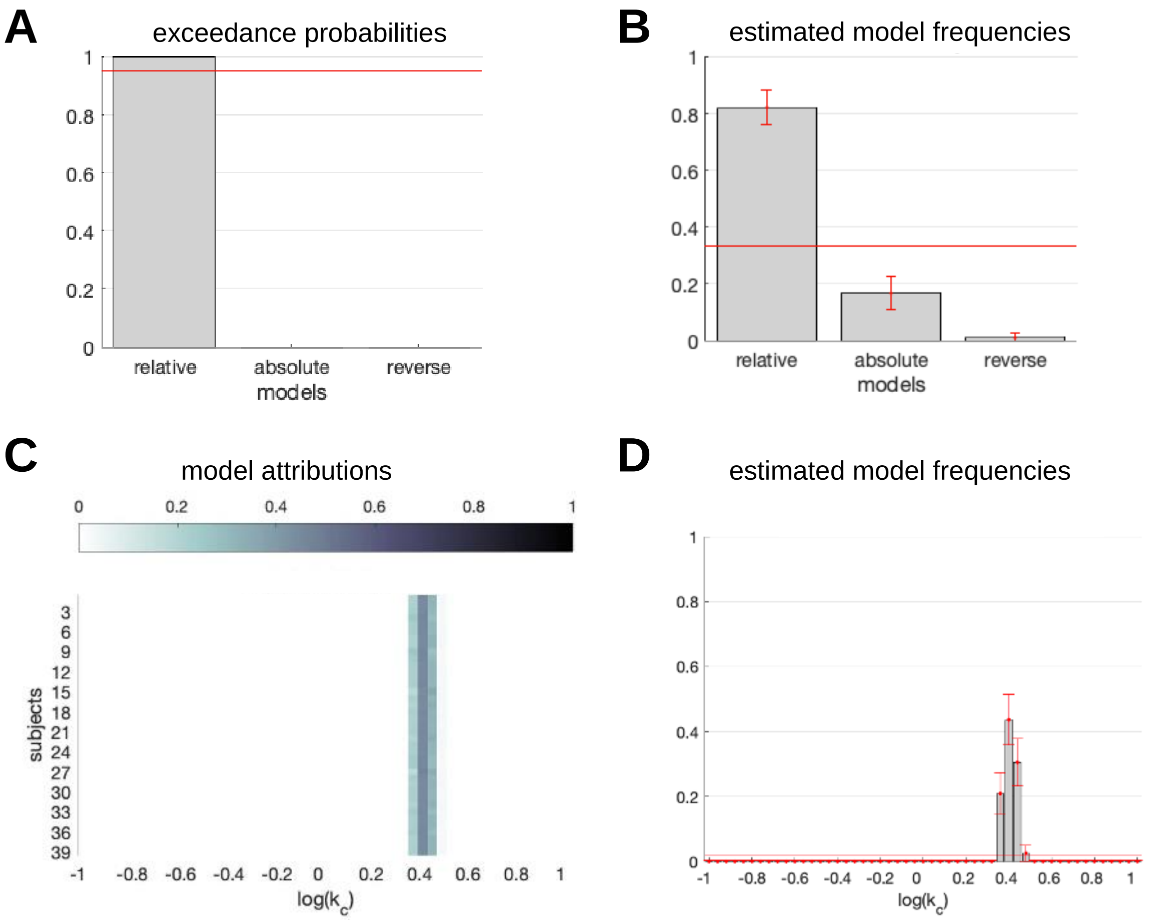


**Figure S5. Bayesian model selection for line normalization ratios (related to Fig. 5).** (A-B) Graphs of the exceedance probabilities and estimated model frequencies for Bayesian Model Selection (BMS) across three models: a 'relative' model in which the neural data of individual participants are fit with 3D lines of equal length, and $\log\left(\frac{16}{11}\right) \sim 0.38$ so that $k_{low} = k_{high} = k_{full}$; an 'absolute' model in which the length of each model line is in proportion to the number of elements within it (the line corresponding to the full range context is longer than the other two, $\log(k_c)$=0); a 'reverse' model in which the lines for the low and high contexts are longer than that of the full context $(\log\left(\frac{11}{16}\right) \sim -0.38)$, a reverse "anti-normalisation". Exceedence probability measure how likely it is that any given model is more frequent than all other models in the set (Stephan et al, 2009). Expected frequencies are the expected number of participants explained by each model. The red horizontal line indicates the null frequency profile across models. (C-D) BMS considering instead the whole range of possible line length ratios. (C) Plot of model attributions, which are an aggregate estimate of how likely it was that each participant was best explained by each of the models (models spanning line length ratios -1 < $\log(k_c)$ < 1). (D) Expected frequencies for the full range of line length ratios. We find strong evidence favoring a model with $\log(k_c) \sim 0.4$, which corresponds more or less exactly to full normalisation. Together, this evidence provides qualitative and quantitative evidence showing that our effects hold at the per-individual level.
